# Quantifying biological heterogeneity in nano-engineered particle-cell interaction experiments

**DOI:** 10.1101/2025.02.01.636020

**Authors:** Ryan J. Murphy, Matthew Faria, James M. Osborne, Stuart T. Johnston

**Affiliations:** UniSA STEM, The University of South Australia, Mawson Lakes, SA 5095, Australia; School of Mathematics and Statistics, The University of Melbourne, Parkville, Victoria, Australia; Department of Biomedical Engineering, The University of Melbourne, Parkville, Victoria, Australia

## Abstract

Nano-engineered particles are a promising tool for medical diagnostics, biomedical imaging, and targeted drug delivery. Fundamental to the assessment of particle performance are *in vitro* particle-cell interaction experiments. These experiments can be summarised with key parameters that facilitate objective comparisons across various cell and particle pairs, such as the particle-cell association rate. Previous studies often focus on point estimates of such parameters and neglect heterogeneity in routine measurements. In this study, we develop an ordinary differential equation-based mechanistic mathematical model that incorporates and exploits the heterogeneity in routine measurements. Connecting this model to data using approximate Bayesian computation parameter inference and prediction tools, we reveal the significant role of heterogeneity in parameters that characterise particle-cell interactions. We then generate predictions for key quantities, such as the time evolution of the number of particles per cell. Finally, by systematically exploring how the choice of experimental time points influences estimates of key quantities, we identify optimal experimental time points that maximise the information that is gained from particle-cell interaction experiments.

## 1 Introduction

Nano-engineered particles have the potential to transform precision medicine [1–4], medical diagnostics and biomedical imaging [5–8]. They facilitate the targeted delivery of therapeutics and imaging agents to specific cell types in challenging biological environments [3, 9, 10]. Particles can be designed and produced with varying physicochemical characteristics tailored to specific applications, such as cancer medicines, gene therapies, immunotherapies [3], and vaccines [11, 12]. However, determining the performance of a particular design for a particular application is challenging. This is because many biological, physical, and chemical processes occur simultaneously across a range of spatial and temporal scales [10, 13–18]. Here, we develop tools to assess the performance of nano-engineered particle designs using a combined mathematical-statistical-experimental framework that quantifies crucial nano-engineered particle-cell interactions [13–16].

*In vitro* particle-cell interaction experiments provide a relatively fast and inexpensive option to build an understanding of nano-engineered particle-cell interactions (Fig. 1). In these experiments, nano-engineered cells are incubated with particles over a period of hours to days [19]. Measurements estimating the number of associated particles per cell can be obtained in different ways, including microscopy, flow cytometry, and spectrometer-based methods. A review of these techniques can be found in [20]. Microscopy techniques provide data at a range of spatial resolutions, including approximate counts of individual nanoparticles at the single cell level, but generating representative samples is time and labour intensive [21–23]. In this study, we focus on routinely generated, rapid, and high-throughput flow cytometry data [19, 24, 25]. Throughout we use the term nano-engineered particles, or *particles* for brevity, to describe nanoparticles (less than 100nm in diameter) and particles that are hundreds of nanometres in diameter generated using nano-engineering techniques [10].

**Figure 1:**
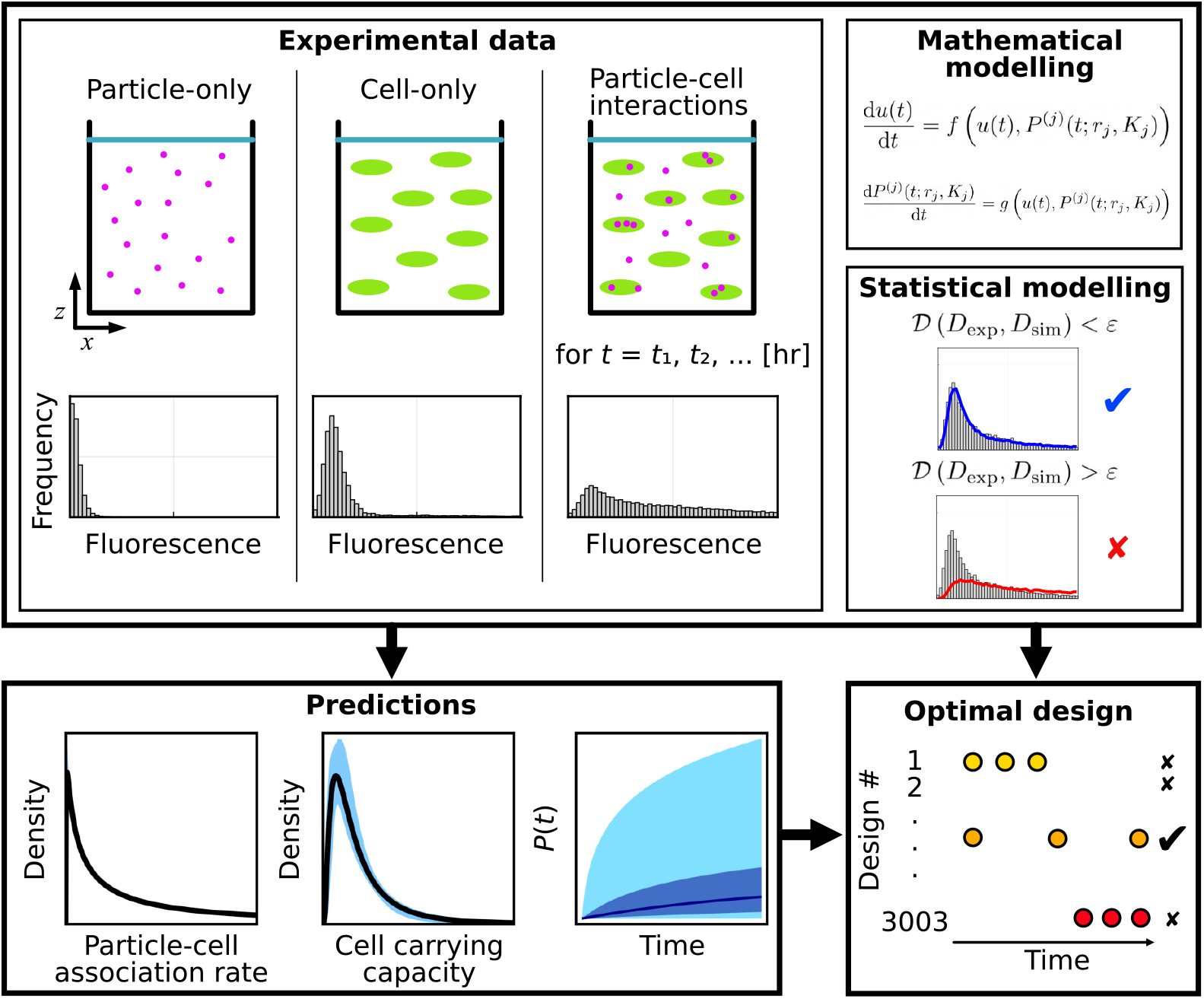
Mathematical-statistical-experimental workflow to analyse particle-cell interaction experiments. Experimental data comprises flow cytometry measurments from particle-only, cell-only, and particle-cell experiments. We use mathematical and statistical modelling to generate predictions of key quantities and identify optimal experimental time points.

Mathematical modelling provides a powerful tool to characterise particle-cell interactions, reviewed in [10, 26], and interpret data from particle-cell interaction experiments. In particular, mechanistic mathematical modelling has been used to reveal key mechanisms driving particle-cell interaction experiments, including particle transport [27–31] and internalisation processes [26, 32–34]. In such studies there is a growing recognition of the importance of heterogeneity in biological and physical processes and properties [35–38]. Previous studies have also provided a mechanistic and quantitative basis for understanding particle-cell interactions using experimentally validated differential equation-based models [19, 35, 38]. These studies have revealed that particle-cell interactions are well characterised by two parameters: *particle-cell association rate*, analogous to a rate constant in chemical reactions, and a *carrying capacity-type*, which characterises the maximum number of particles associated with a cell. This approach groups together particles in contact with the surface of a cell and particles internalised within a cell [19]. Using such metrics has been recommended as a route to accelerate progress by enabling objective comparisons across data generated using different experimental protocols or pairs of cells and particles [39–41].

In this study, we generalise the mechanistic ordinary differential equation-based mathematical model in [19] to biologically heterogeneous cell populations. Specifically, we allow the particle-cell association rate, carrying capacity, and autofluorescence to be distinct for each cell and allow the fluorescence of each particle to be distinct. This new mathematical model captures particle-cell interactions in experiments with cell-cell heterogeneity and captures cell-cell competition for particles. To perform parameter estimation, practical identifiability analysis, and prediction for the mathematical model, we use approximate Bayesian computation methods that have been developed to interpret similar flow cytometry data [32, 43–46]. These approximate Bayesian computation methods seek parameter values that minimise the difference between simulated data from the model and observed experimental data. This combined mathematical-statistical-experimental approach allows us to estimate key parameters, variation in key parameters, and uncertainty in key parameters. We directly compare this new model to previous approaches that do not allow for heterogeneous populations, neglect heterogeneity in routine measurements, and apply data transformations that account for heterogeneity in routine measurements but can generate non-physical quantities. This study builds on the growing literature that has focused on quantifying cell-cell heterogeneity from flow cytometry data in the absence of particles [47–53]. We conclude by exploring how the choice of experimental time points influences information gained about the mathematical model parameters from our approximate Bayesian computation approach. In this process, using established methods based on principles of Bayesian optimal design [54–56], we systematically examine 3003 experimental designs, identify optimal designs for different particle-cell association rates, and identify an overall optimal design for when the particle-cell association rate is unknown.

## 2 Data

We examine previously published data for THP-1 cells (a human leukaemia monocytic suspension cell line) separately incubated with three types of nano-engineered particles [19]. Results in the main manuscript focus on 150 nm polymethacrylic acid(PMA) core-shell particles. Supplementary results focus on 214 nm PMA-capsules and 633 nm PMA core-shell particles.

Data for each particle-cell combination are obtained by flow cytometry and comprise of (i) 20,000 measurements per time point of the fluorescent signal for individual cells in time course particle-cell association data, 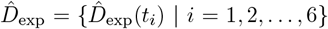 for *t*_*i*_ = 1, 2, 4, 8, 16, and 24 [hr] (Fig. 1(C,F)); (ii) 11,605 measurements of the fluorescent signal for individual cells for the cell-only control data, 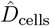, corresponding to measurements at *t* = 0 [hr] (Fig. 1(B,E)); and, (iii) at least 249,344 measurements of fluorescent signal for individual particles for a particle-only control data, 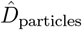 (500,000 measurements for 150 nm PMA core-shell particles and 214 nm PMA-capsules, 249,344 measurements for 633 nm PMA core-shell particles) (Fig. 1(A,D)).

Following [19], these measurements are calibrated to a reference voltage and denoted *D*_exp_, *D*_cells_, and *D*_particles_ (Supp. S1.1). Data for each time point group together endpoint measurements from two replicates. We do not obtain multiple measurements of the same cell. These types of data are sometimes referred to as snapshot time-series data [49].

## 3. Methods

We seek to (i) quantify biological heterogeneity in nano-engineered particle-cell experiments, (ii) generate predictions of key quantities that are challenging to observe by experimentation alone, and (iii) identify experimental time points that are optimal in the sense of maximising the precision of parameter estimates. To perform this analysis we present a suite of quantitative techniques: two mechanistic mathematical models that describe particle-cell interactions; simulation-based approximate Bayesian computation (ABC) methods that we employ for parameter inference, practical identifiability analysis, and prediction; and experimental design tools based on principles of Bayesian optimal design.

### 3.1. Mathematical models

We consider two ordinary differential equation-based mathematical models. The first is a previously published model that assumes a homogeneous cell population [19]. The second is a new mathematical model that generalises the homogeneous model to heterogeneous cell populations.

#### 3.1.1. Homogeneous cell population

Assuming all cells in the experimental well are identical, the concentration of particles per cell in the well-mixed media of volume *V* at time *t*, denoted *u*(*t*) [particles cell^*−*1^ m^*−*3^], is initially given by *u*(0) = *u*_0_ [particles cell^*−*1^ m^*−*3^] and its time evolution is governed by the ordinary differential equation [19],

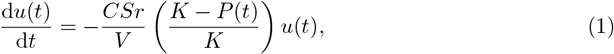

where *C* [-] is the fractional surface coverage of cells, *S* [m^2^] is the surface area of the cell boundary, *r* [m s^*−*1^] is the particle-cell association rate, and we refer to *K* [particles cell^*−*1^] as the cell carrying capacity for particles. In practice, Eq. (1) is a phenomenological model and particles may rapidly associate and disassociate, so *K* is not necessarily a carrying capacity in the traditional sense of population dynamics models, however it captures the observed behaviour. Considering conservation of the total number of particles in the system and assuming that there is no particle degradation during the experiment, the number of associated particles per cell at time *t*, denoted *P* (*t*) [particles cell^*−*1^], is equal to the difference between the initial number of particles per cell in the media and the number of particles per cell in the media at time *t*,

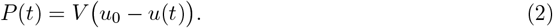

The analytical solution of the coupled system of Eqs. (1)-(2) for *P* (*t*) is,

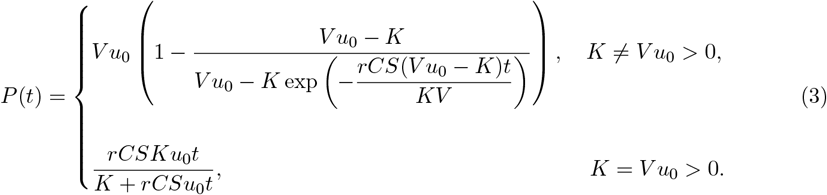

Whether the solution approaches the long-time solution, *P* (*t*) = *K*, within the experimental duration depends on *r*. In typical experiments, the number of particles per cell is small relative to the initial number of particles in the media, *P* (*t*) ≪ *V u*_0_ for all *t* [19].

This mechanistic mathematical model is characterised by four parameters (*C, S, V, u*_0_), that are directly measured in the experiments and assumed to be known fixed constants (Supp. S1.2), and two unknown parameters, (*r, K*), that cannot be directly measured in the experiments.

#### 3.1.2. Heterogeneous cell population

Previous studies have shown that the particle-cell association rate, *r*, and the carrying capacity parameter, *K*, exhibit significant variability between cells subjected to the same experimental conditions [35]. We now generalise the homogeneous cell population model to allow for this variability. We make a common assumption for non-negative biological parameters and assume that *r* and *K* are lognormally distributed [27, 57, 58],

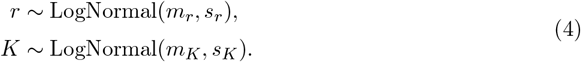

In Eq. (4), *m*_*r*_ *>* 0 and *m*_*K*_ *>* 0 denote the means and *s*_*r*_ *>* 0 and *s*_*K*_ *>* 0 denote the standard deviations of the lognormal distributions for *r* and *K*, respectively. We use this parameterisation of the lognormal distributions to report the results for *r* and *K* in terms of their mean and standard deviation. With this approach, we allow for differing amounts of heterogeneity in the particle-cell association rate, *r*, and the carrying capacity, *K*, and assume that there is no correlation between these mechanisms for each individual cell [35].

Assuming that each cell has distinct properties, the time evolution of the concentration of particles per cell in the well-mixed media can be described by a system of *Ñ* + 1 ordinary differential equations, where *Ñ* is the number of cells simulated in the experimental well. We set *Ñ* = 20, 000, which is equal to the number of experimental measurements per time point and assume that this is sufficiently large to accurately approximate the *r*-*K* distribution. This model captures cell-cell competition for particles. For example, if a population of cells all have similar values of *K*, those cells with high *r* reduce the total number of particles available to associate with cells with low *r* at later times. In Supp. S1.3, we derive this heterogeneous model and show that the heterogeneous model simplifies to the homogeneous model when all cells are identical.

Solving the large system of coupled differential equations that forms the heterogeneous model is computationally expensive, and typically takes minutes to solve. This makes statistical inference, where we must solve the model many times for different parameter values, computationally challenging. To address this challenge, we determine an approximate solution to the heterogeneous model that is accurate under the experimentally relevant assumption that the number of particles that associate with cells is small relative to the initial number of particles in the media. Furthermore, this approximate solution can be evaluated in less than a second, which supports efficient statistical inference. The approximate solution to the heterogeneous model, derived in Supp. S1.3, is given by a set of independent equations for the number of particles associated with each cell *j* = 1, 2, … *Ñ*,

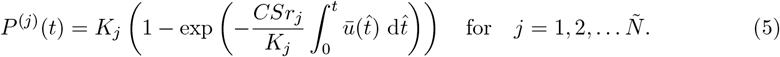

In Eq. (5), *r*_*j*_ and *K*_*j*_ are samples from the probability distributions defined in Eq. (4) and *ū*(*t*) [particles cell^*−*1^ m^*−*3^] represents the approximate concentration of particles per cell in the media at time *t*. We estimate *ū*(*t*) using the mean of *M* evaluations of the analytic solution to the homogeneous cell population model (Eq. (3)) at distinct samples of *r* and *K* from Eq. (4). We set *M* = 100 based on pilot simulations exploring a tradeoff between computational efficiency and accuracy. We evaluate the integral in Eq. (5) using the trapezoid rule. In Supp. S1.3, we verify that this approximate solution to the heterogeneous model (Eq. (5)) agrees with the corresponding solution to the heterogeneous model for experimentally relevant parameter regimes.

This mechanistic mathematical model is characterised by four parameters, (*C, S, V, u*_0_), that are directly measured in the experiments and assumed to be known fixed constants (Supp. S1.2), and four unknown hyperparameters, (*m*_*r*_, *s*_*r*_, *m*_*K*_, *s*_*K*_), that we will estimate as they cannot be directly measured in the experiments.

### 3.2. Parameter estimation, practical identifiability analysis, and prediction

We use established likelihood-free simulation-based ABC methods for parameter estimation, practical identifiability analysis, and prediction. In brief, we use ABC methods that seek parameter values that minimise the difference between the experimentally measured flow cytometry fluorescence data and synthetic flow cytometry data that we generate by simulation.

For illustrative purposes, we first explain how to generate synthetic flow cytometry data under simplifying assumptions. In particular, we assume that the homogeneous mathematical model is valid and that it is reasonable to summarise the experimentally measured cell-only and particle-only control data sets, *D*_cells_ and *D*_particles_ defined in Sec. 2, with their median fluorescent intensities. The only synthetic flow cytometry datapoint at time *t*_*i*_ is then [19, 41]

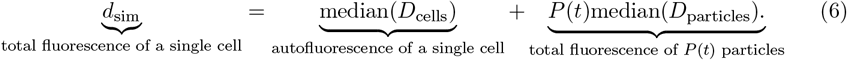

In Eq. (6), the contribution of cell autofluorescence to the total fluorescence is additive, and the contribution of particle fluorescence to the total fluorescence is multiplicative.

In practice, we generalise Eq. (6) to allow for the heterogeneous outputs from the heterogeneous mathematical model and we capture cell-cell variability, particle-particle variability, and measurement noise by exploiting heterogeneity in *D*_cells_ and *D*_particles_. A single flow cytometry datapoint for cell *j* = 1, 2, …, 20000 at time *t*_*i*_ is then given by,

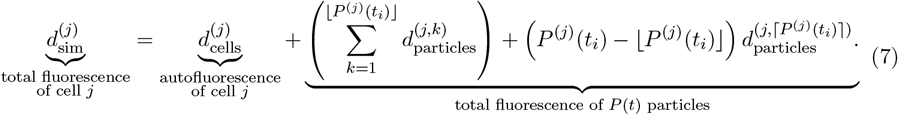

Analogous to Eq. (6), in Eq. (7) noise due to cell autofluorescence, ^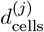^, is additive but rather than using the median fluorescent intensity of *D*_cells_, we sample the autofluorescence _of each cell *j*,_ ^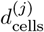^, from *D*_cells_. Similarly, we refer to noise due to the particle fluorescence as multiplicative, but rather than using the median fluorescent intensity, we now sample the fluorescence of each individual particle *k* for cell *j*, ^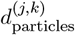^, from *D*_particles_. In Eq. (7), since *P* ^(*j*)^(*t*_*i*_) may not be an integer, we assume that the total fluorescence due to particles is given by ⌊*P* ^(*j*)^(*t*_*i*_)⌋ samples from *D*_particles_, where ⌊·⌋ denotes the floor function, and one additional sample from *D*_particles_ that is scaled by the fraction of the particle that is associated with the cell, ^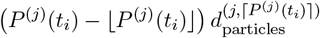^, where ⌈·⌉ denotes the ceiling function.

A complete synthetic flow cytometry time course data set is then *D*_sim_ = {*D*_sim_(*t*_*i*_) | *t*_*i*_ = 1, 2, 4, 8, 16, 24}, where 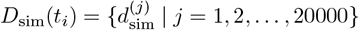. As we simulate the synthetic flow cytometry datapoints at each time point *t*_*i*_ and for *j* = 1, 2, …, 20000 we extensively sample *D*_cells_ and *D*_particles_. Throughout, we use capitalised *D* and non-capitalised *d* to distinguish between data sets and the data points that form data sets, respectively.

We perform inference using an established ABC-Sequential Monte Carlo (SMC) algorithm [32, 46] that we modify to terminate at a target ABC error threshold. To facilitate efficient inference, we use an ABC-SMC algorithm, which sequentially determines sets of parameter values that result in closer agreement between the synthetic and experimental time course data sets, *D*_sim_ (defined above) and *D*_exp_ (defined in Sec. 2), respectively. We use uniform priors for all parameters and set the ABC distance function to be the sum of the Anderson-Darling distance between the experimental and simulated datasets at each time point. In Supp. S1.4, we present further details, including the ABC error thresholds, the number of ABC particles, the transition kernel, definitions of the ABC distance functions that we explore, uniform prior bounds, and how we generate posterior distributions, predictions, and inferred distributions.

### 3.3. Optimal experimental time points

To identify optimal experimental time points, we adopt established Bayesian optimal design techniques for mathematical models with intractable likelihoods. Specifically, we consider each set of time points as a distinct experimental design *w* and take an approach similar to the ABCdE algorithm [54] that involves ABC rejection and pre-simulating synthetic data.

For consistency with previous data [19], we assume that each experimental design comprises flow cytometry measurements at six distinct time points in addition to cell-only control data (corresponding to *t* = 0 [hr]) and particle-only control data. For each particle-cell scenario that we consider we assume that the cell-only and particle-only control data are fixed. Each experimental design is then characterised by six distinct time points. We assume that these time points are chosen from fourteen possible choices: *t* = 0.5 [hr], *t* = 1 [hr] and every two hours from *t* = 2 [hr] to *t* = 24 [hr]. These time points are chosen based on previous data and practical considerations regarding the frequency of measurements with manual experimental procedures. As we consider all possible ways to choose the six time points from a set of fourteen possible time points, we examine 3003 possible experimental designs.

Given that particles are frequently designed to target specific cells, the particle-cell association rate *r* is a key quantity of interest. As the particle cell association rate *r* has been shown to vary over multiple orders of magnitude for different combinations of particles and cell types [19], we examine each of the 3003 experimental designs for three particle-cell scenarios that vary with respect to the particle-cell association rate *r*. We refer to these as low *r*, intermediate *r*, and high *r* and note that these are defined relative to *K*.

For each scenario, we capture potential variation in experimental data by pre-simulating *J* = 20 synthetic flow cytometry data sets using statistical hyperparameters *m*_*r*_, *s*_*r*_, *m*_*K*_, *s*_*K*_ sampled from non-negative truncated Gaussian distributions with pre-specified mean values and coefficients of variation set equal to 0.1. We next pre-simulate *N*_pre_ = 200, 000 synthetic flow cytometry data sets using statistical hyperparameters sampled from uniform priors whose bounds are defined in Supp. S1.4. Using ABC rejection across all designs, we use these *N*_pre_ data sets to efficiently form ABC posteriors. We choose the ABC error threshold so that all ABC posteriors across all designs contain at least 200 samples and use the Anderson-Darling ABC distance throughout.

To assess each design *w* for a particular scenario we compute an average utility that rewards precise estimates of the model parameters *θ* = (*m*_*r*_, *s*_*r*_, *m*_*K*_, *s*_*K*_) across the *J* synthetic flow cytometry data sets via the empirical covariance matrix of the ABC posterior distributions [55],

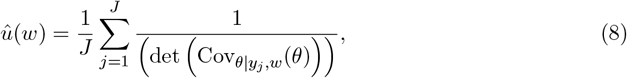

In Eq. (8), *y*_*j*_ represents a single synthetic flow cytometry data set, and ABC posteriors are computed for each of the *J* synthetic data sets independently.

The optimal design for a particular scenario corresponds to the design that maximises the average utility over all of the 3003 possible designs, *W*, with respect to potential future data and model parameters,

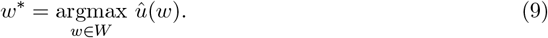

## 4 Results and discussion

Particle-cell interaction experiments generate thousands of cell-level measurements per time point. For example, we analyse experiments with 20,000 measurements at each time point [19]. However, heterogeneity that is present in such data is often overlooked when using standard metrics such as the median fluorescence intensity. Here, we use mechanistic mathematical modelling and statistical parameter estimation, practical identifiability analysis, and prediction tools to exploit this heterogeneity in routine measurements for greater understanding.

We first perform synthetic data studies to verify that our methods recover known parameters and quantities, explore the role of heterogeneity on key quantities, and to demonstrate parameter identifiability challenges that may arise. We then apply these methods to experimental data to quantify biological heterogeneity, quantify uncertainty in estimates of biological heterogeneity, and generate predictions of key quantities that are challenging to determine by experimentation alone. Next, we compare our new approach to a previous method. We conclude by identifying optimal experimental time points across various particle-cell scenarios.

### 4.1. Mechanistic modelling incorporating biological and control data heterogeneity captures data variability and reveals sources of uncertainty

Before analysing experimental data we verify that the parameter inference methods that we employ recover known parameters and quantities. We generate synthetic flow cytometry data consistent with the experimental data (Section 3). Heterogeneity in this synthetic data is driven by prescribed biological heterogeneity and heterogeneity in routine control experimental data measurements. The prescribed biological heterogeneity describes cell-cell variability in particle-cell association rates, *r*, and carrying capacities, *K*. Heterogeneity from the the cell-only and particle-only control experimental data arises in the synthetic data as additive noise due to cell autofluorescence and multiplicative noise due to the presence of particles, respectively.

Analysing this synthetic flow cytometry data using the model that generated the data is useful to explore potential sources of uncertainty in the absence of model misspecification. However, the synthetic data is finite, non-ideal, incomplete, and noisy. Furthermore, while it is straightforward to simulate the model, which incorporates biological heterogeneity and heterogeneity from the control experimental data, the complexity of the model renders traditional likelihood-based inference techniques computationally challenging.

Using likelihood-free simulation-based ABC inference methods (Section 3.2), we form posterior distributions for the means and standard deviations of the lognormal distributions that we assume characterise *r* and *K*. We use these posterior distributions to capture uncertainty and estimate credible intervals, which we report in terms of highest posterior density. For these synthetic data, we find that posterior distributions for all statistical hyperparameters are relatively narrow in comparison to pre-specified bounds and are well-formed about a single peak. Therefore, we conclude that the statistical hyperparameters (*m*_*r*_, *s*_*r*_, *m*_*K*_, *s*_*K*_) are practically identifiable (Fig. 2). This means that a relatively narrow range of parameters gives similar agreement to the experimental data. Furthermore, the known values of the statistical hyper-parameters used to generate the data are each contained within their respective 95% highest posterior density interval.

**Figure 2:**
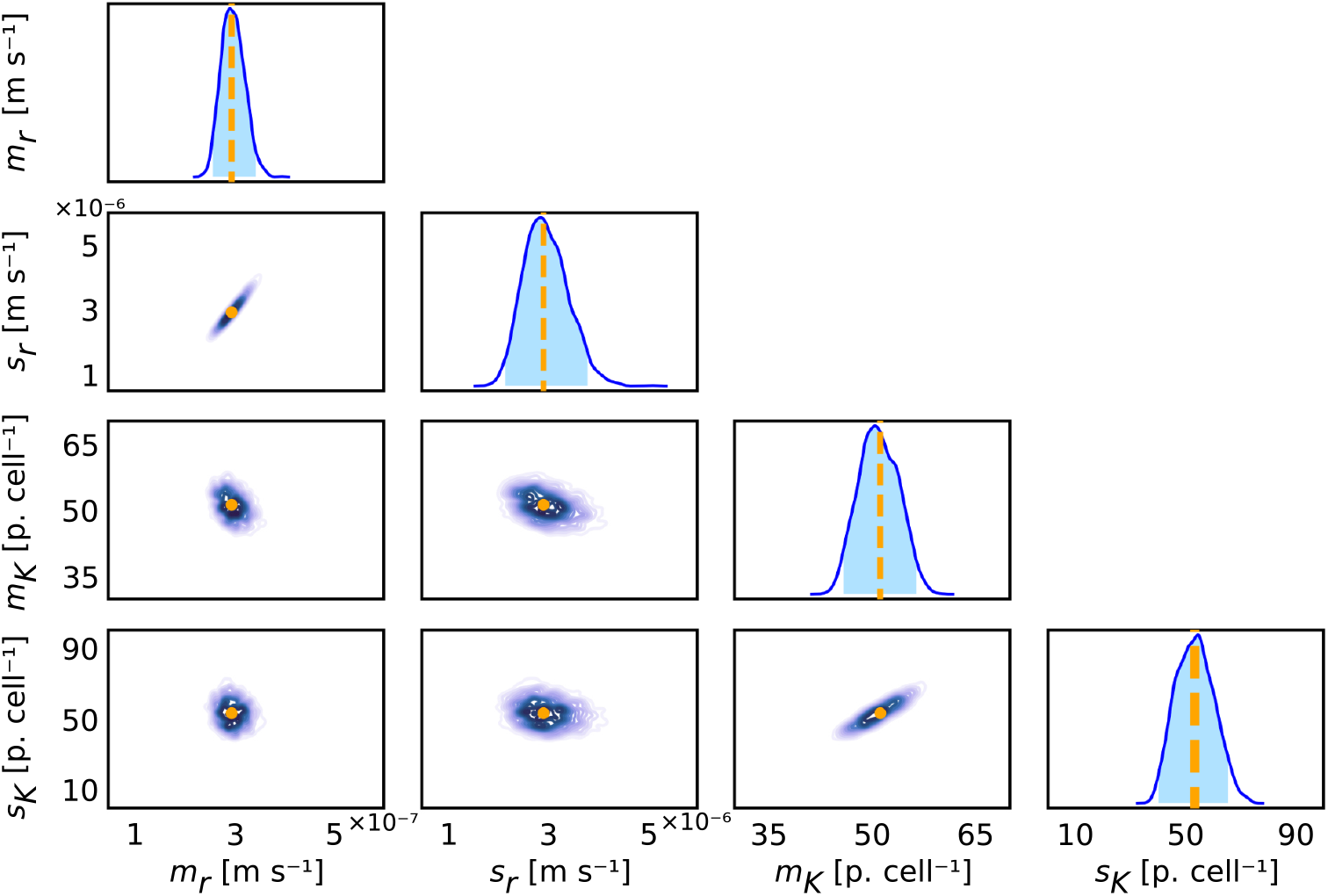
Parameter inference techniques recover known values of the statistical hyperparameters from finite, non-ideal, incomplete, and noisy synthetic data. Univariate and bivariate posterior distributions for the statistical hyperparameters *m*_*r*_ [m s^*−*1^], *m*_*K*_ [particles cell^*−*1^], *s*_*r*_ [m s^*−*1^], and *s*_*K*_ [particles cell^*−*1^]. The 95% univariate highest posterior density intervals are (2.58 × 10^*−*7^, 3.44 × 10^*−*7^) for *m*_*r*_, (2.16 × 10^*−*6^, 3.81 × 10^*−*6^) for *s*_*r*_, (45.8, 56.4) for *m*_*K*_, and (40.1, 65.3) for *s*_*K*_. Known parameter values used to generate the data are (*m*_*r*_, *s*_*r*_, *m*_*K*_, *s*_*K*_) = (2.96 × 10^*−*7^, 2.92 × 10^*−*6^, 51.2, 53.3) and are shown as vertical orange dashed lines and circles.

Propagating forward the uncertainty captured by the posterior distributions is a powerful technique for additional verification and allows for comparisons to experimentally measured quantities. We use this technique to show that posterior predictions for the distribution of fluorescence measurements demonstrate excellent agreement with the corresponding synthetic data (Fig. 3).

**Figure 3:**
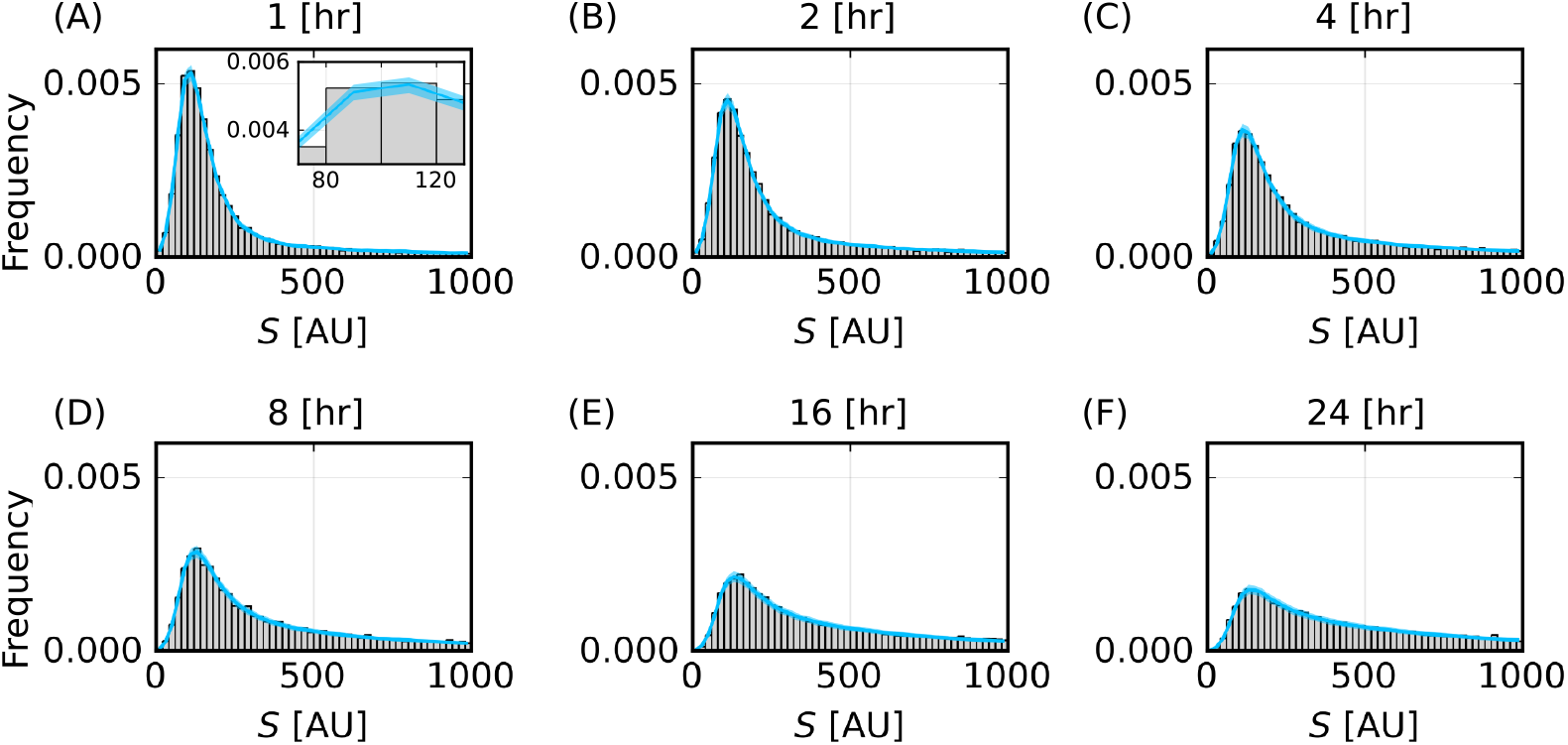
Posterior prediction of particle-cell association time course data captures variability in the synthetic data. Histograms of the synthetic fluorescence signal, *S* [AU], (grey) and 95% prediction interval of heights of histogram bars from the mathematical model (cyan) for (A) *t* = 1, (B) 2, (C) 4, (D) 8, (E) 16, (F) 24 [hr]. Inset in (A) illustrates the width of prediction intervals.

We next propagate forward uncertainty captured by the posterior distributions for the statistical hyperparameters to generate inferred distributions for *r* and *K* (Fig. 4). These inferred distributions allow us to quantify biological heterogeneity and allow us to quantify the uncertainty in these estimates of the biological heterogeneity. These inferred distributions demonstrate that there is significant biological heterogeneity. Previous methods that focus on point estimates of parameters that characterise particle-cell interactions in the homogeneous model cannot capture this heterogeneity [19]. Furthermore, while more recent methods can capture biological heterogeneity, they overlook heterogeneity that is present in routine control experimental data measurements that we later show is critical for generating accurate predictions [35].

**Figure 4:**
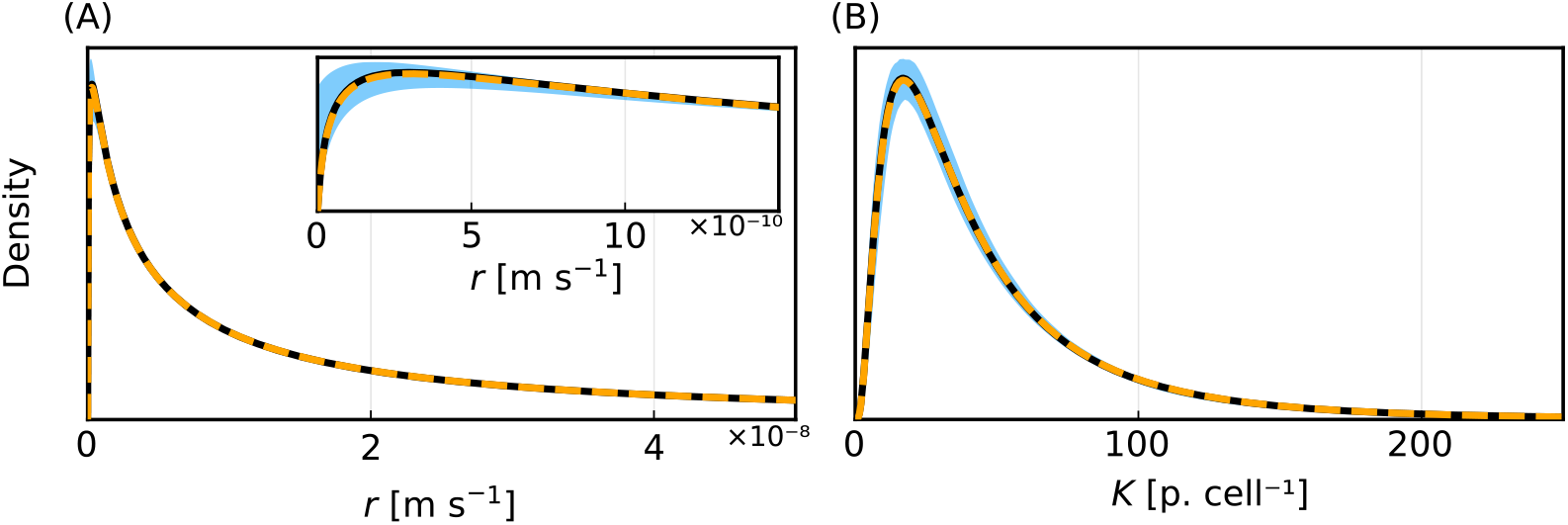
Parameter inference techniques recover known inferred distributions from finite, non-ideal, incomplete, and noisy synthetic data. (A) Inferred distribution for *r* [m s^*−*1^] with inset focusing on low *r*. (B) Inferred distribution for *K* [particles cell^*−*1^]. Colours represent median of estimated inferred distribution (black), 95% prediction interval (blue), and distributions at known values (orange-dashed).

### 4.2. Mathematical modelling enables predictions of key quantities that are challenging to observe experimentally

Particle-cell interaction experiments are routinely performed to address a fundamental question: how does the number of particles per cell change with time? This is a challenging question to answer accurately and robustly by experimentation alone. Counting the number of particles per cell at scale in experiments is simply not yet feasible. Furthermore, straightforward translations of fluorescence measurements to particles per cell, such as in [19], assume homogeneous cell populations and do not generate robust accurate estimates for the number of particles per cell, as we later show. Our mathematical and statistical modelling framework provides a powerful tool to address this question. Propagating forward the uncertainty in posterior distributions, we generate predictions for the time evolution of the number of particles per cell, *P* (*t*). These predictions demonstrate excellent agreement with the corresponding synthetic data. In particular, the lower and upper quartiles of the known synthetic data demonstrate excellent agreement with the corresponding boundaries of the 50% prediction interval (Fig. 5(A)). This is a critical verification step for our approach. This prediction for *P* (*t*) is obtained by analysing the synthetic flow cytometry data that comprises noisy observations of *P* (*t*) as described in Eq. (7), whereas the synthetic data that we make a comparison to are noise-free observations of *P* (*t*) obtained from Eq. (5).

**Figure 5:**
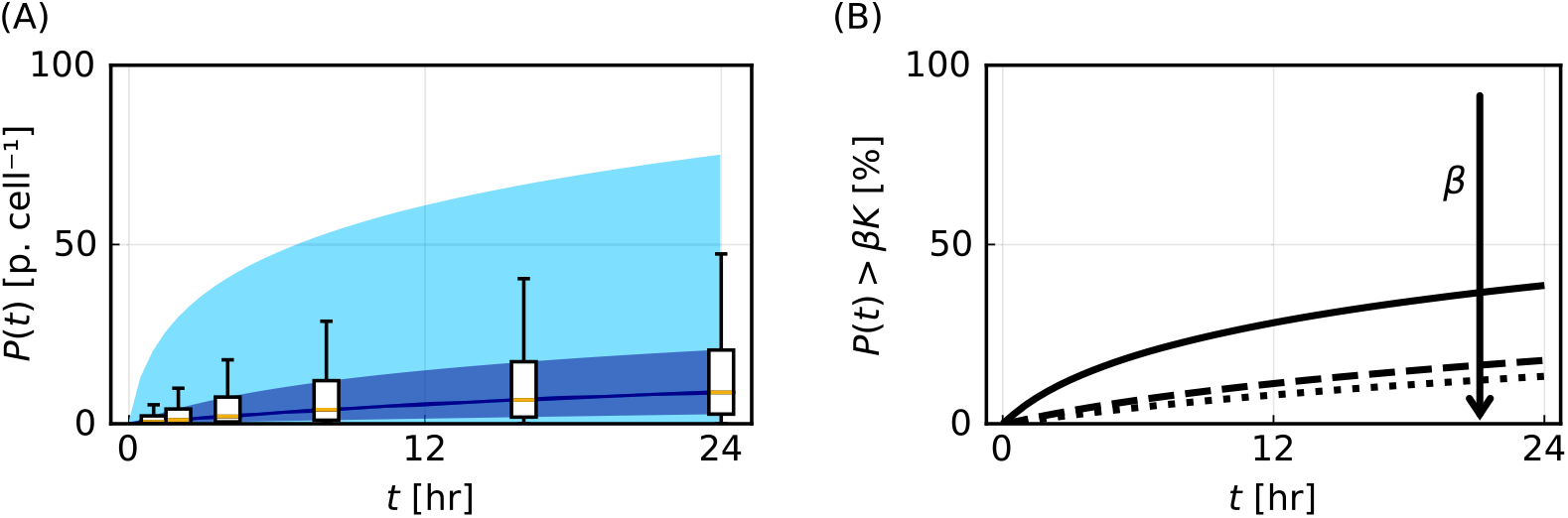
Mathematical modelling generates predictions of quantities that are challenging to observe experimentally. (A) Time evolution of the number of particles per cell, *P* (*t*) [particles cell^*−*1^], with 50% (dark blue) and 95% (light blue) prediction intervals. Box plots represent synthetic data (outliers not shown). (B) Time evolution of the percentage of cells that are close to carrying capacity, where close is defined via *P* (*t*) *> βK*, for *β* = 0.50 (solid), 0.95 (dashed), and 0.99 (dotted). The arrow indicates the direction of increasing *β*.

We can also use our modelling framework to generate predictions for other quantities that are challenging to observe by experimentation and previous modelling approaches that assume a homogeneous cell population. For example, we can predict the percentage of cells close to their respective carrying capacities (Fig. 5(B)). This could be used to optimise the particle dosage. If most cells are far from the respective carrying capacities increasing the particle dosage could help to elucidate particle-cell interactions. In contrast, if most cells are close to the respective carrying capacities increasing the dosage provides similar results with increased wastage of particles. Predicting the percentage of cells close to carrying capacity can also help interpret uncertainty in estimates of *K*, since parameter estimates for *K* improve when cells are close to their respective carrying capacities.

Our predictions also extend beyond previous flow cytometry-based analyses that classify a cell as being either positive or negative for the presence of particles based on a cutoff fluorescence intensity [25]. We can estimate the percentage of cells close to their respective carrying capacities and the percentage of cells where *P* (*t*) is above a certain threshold. Such cell-level information could be used to identify if particles associate with cells too slowly or too quickly, which may result in subtherapeutic dosages for a particular application or toxic side effects.

### 4.3. Regimes with parameter identifiability challenges

Point estimates for the particle-cell association rate, *r*, have been shown to vary over multiple orders of magnitude for different combinations of particles and cell types [19]. In Figs. 2-5 we consider a scenario where we obtain multiple measurements of the initial increase in *P* (*t*) far from *K* and multiple measurements when *P* (*t*) is close to *K* (Fig. 5(A)). We refer to this as a scenario with intermediate *r*. We now compare a scenario with intermediate *r* to a scenario with low *r*, which only comprises measurements of the initial increase in *P* (*t*), far from *K*, and to a scenario with high *r*, which only comprises measurements of *P* (*t*) close to *K*. Note that these classifications that we base on *r* are relative to *K*. Furthermore, the classifications are based on a prediction of *P* (*t*) that we obtain using our modelling approach.

Performing additional synthetic data studies we analyse how results differ for low, intermediate, and high *r*. We explore these regimes by varying *m*_*r*_ while holding the coefficient of variation *s*_*r*_*/m*_*r*_, the lognormal distribution for *K*, and time points fixed. Exemplar synthetic data for *P* (*t*) simulated for low, intermediate, and high values of *r* are shown in Fig. 6(M-O).

**Figure 6:**
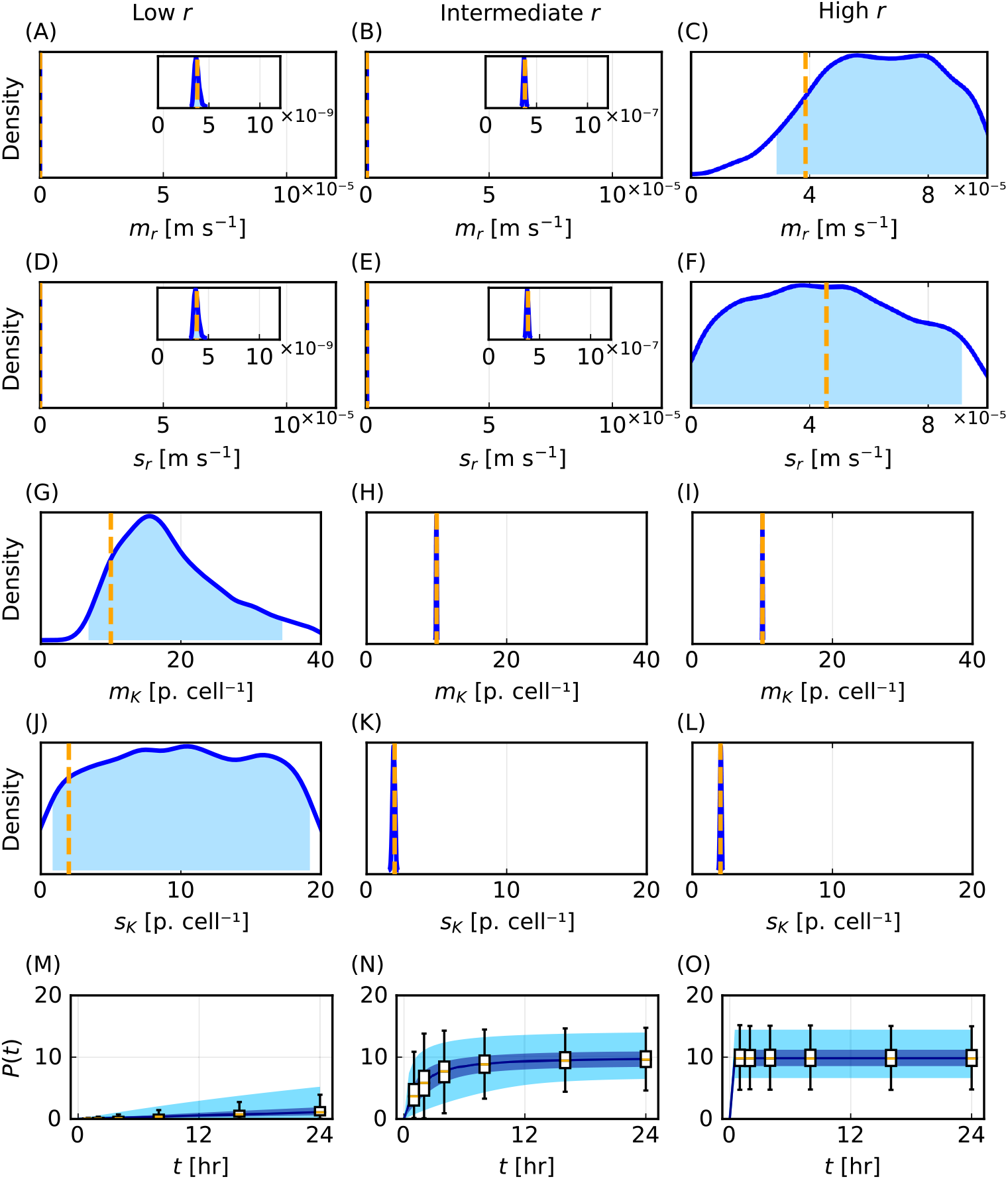
Parameter identifiability challenges for low and high. *r*. Univariate posterior distributions for the statistical hyperparameters *m*_*r*_ [m s^*−*1^], *m*_*K*_ [particles cell^*−*1^], *s*_*r*_ [m s^*−*1^], and *s*_*K*_ [particles cell^*−*1^] for (A,D,G,J) low *r*, (B,E,H,K) intermediate *r*, and (C,F,I,L) high *r*.

For low *r*, posterior distributions for *m*_*r*_ and *s*_*r*_ are relatively narrow in comparison to the pre-specified bounds and are well-formed about a single peak, so we conclude that these parameters are practically identifiable (Fig. 6(A,D)). These parameters are identifiable because we obtain multiple measurements when the influence of *r* is prominent. In contrast, posterior distributions for *m*_*K*_ and *s*_*K*_ are wide and flat relative to pre-specified bounds (Fig. 6(G,J)), so we conclude that these parameters are practically non-identifiable. This lack of identifiability arises because we do not obtain measurements when *P* (*t*) is close to *K*.

For high *r* we obtain results that are opposite to the results for low *r*. We find that *m*_*r*_ and *s*_*r*_ are practically non-identifiable (Fig. 6(C,F)) while *m*_*K*_ and *s*_*K*_ are practically identifiable (Fig. 6(I,L)). This is because we do not obtain measurements when *r* has a significant influence, but we do obtain measurements when *P* (*t*) is close to *K*. For intermediate values of *r*, in agreement with results in Figs. 2-5, all parameters are practically identifiable as we obtain measurements when *r* has a significant influence and when *P* (*t*) is close to *K* (Fig. 6(B,E,H,K)).

These results, which demonstrate that model parameters are partially identifiable for low and high *r*, are consistent with observations for homogeneous population models experiencing sigmoidal growth dynamics [59, 60]. As we later discuss, uncertainty in the posterior distributions result in similar uncertainty for inferred distributions for *r* and *K* and previous methods that focus only on point estimates cannot provide such insights regarding the range of parameters consistent with the data.

Posterior predictions for the flow cytometry data for all values of *r* demonstrate close agreement with the corresponding synthetic data despite the lack of parameter identifiability for low and high values of *r* (Supp. Fig. S8). This predictive capability in the presence of poor identifiability is expected because our ABC method involves identifying parameter values that result in close agreement with the synthetic flow cytometry data. This result is consistent with observations for homogeneous population models [61]. As our ABC method is designed only to minimise the difference to flow cytometry data, predictions for other quantities should be explored on a case-by-case basis. In this regard, predictions of *P* (*t*) are found to demonstrate excellent agreement with the corresponding synthetic data for all values of *r* (Fig. 6(M-O)).

Time evolution of the number of particles per cell, *P* (*t*) [particles cell^*−*1^], with 50% (dark blue) and 95% (blue) prediction intervals with box plots representing synthetic data with outliers not shown for (M) low *r*, (N) intermediate *r*, and (O) high *r*. Synthetic data are generated using the following parameter values: (*m*_*K*_, *s*_*K*_) = (10.0, 2.0) for low, intermediate, and high *r* with low *r* (*m*_*r*_, *s*_*r*_) = (3.86 × 10^*−*8^, 4.57 × 10^*−*8^), intermediate *r* (*m*_*r*_, *s*_*r*_) = (3.86 × 10^*−*7^, 4.57 × 10^*−*7^), high *r* (*m*_*r*_, *s*_*r*_) = (3.86 × 10^*−*5^, 4.57 × 10^*−*5^).

### 4.4. Insights from experimental data

We now apply our verified mathematical and statistical modelling tools to experimental data. Working directly with the experimental data poses new challenges as we do not know the true data-generating process. We allow for model misspecification through an increased ABC error threshold compared to synthetic data.

We focus on experiments where 150 nm PMA core-shell particles are incubated with THP-1 cells. Posterior distributions for each statistical hyperparameter suggest that they are practically identifiable (Fig. 7(A-D)). The posterior distributions for *s*_*r*_ and *s*_*K*_ and the inferred distributions for *r* and *K* (Fig. 7(C,D)) both indicate significant cell-cell variability in the population. We also observe significant cell-cell variability in predictions of the number of particles per cell, *P* (*t*) (Fig. 7(E)).

**Figure 7:**
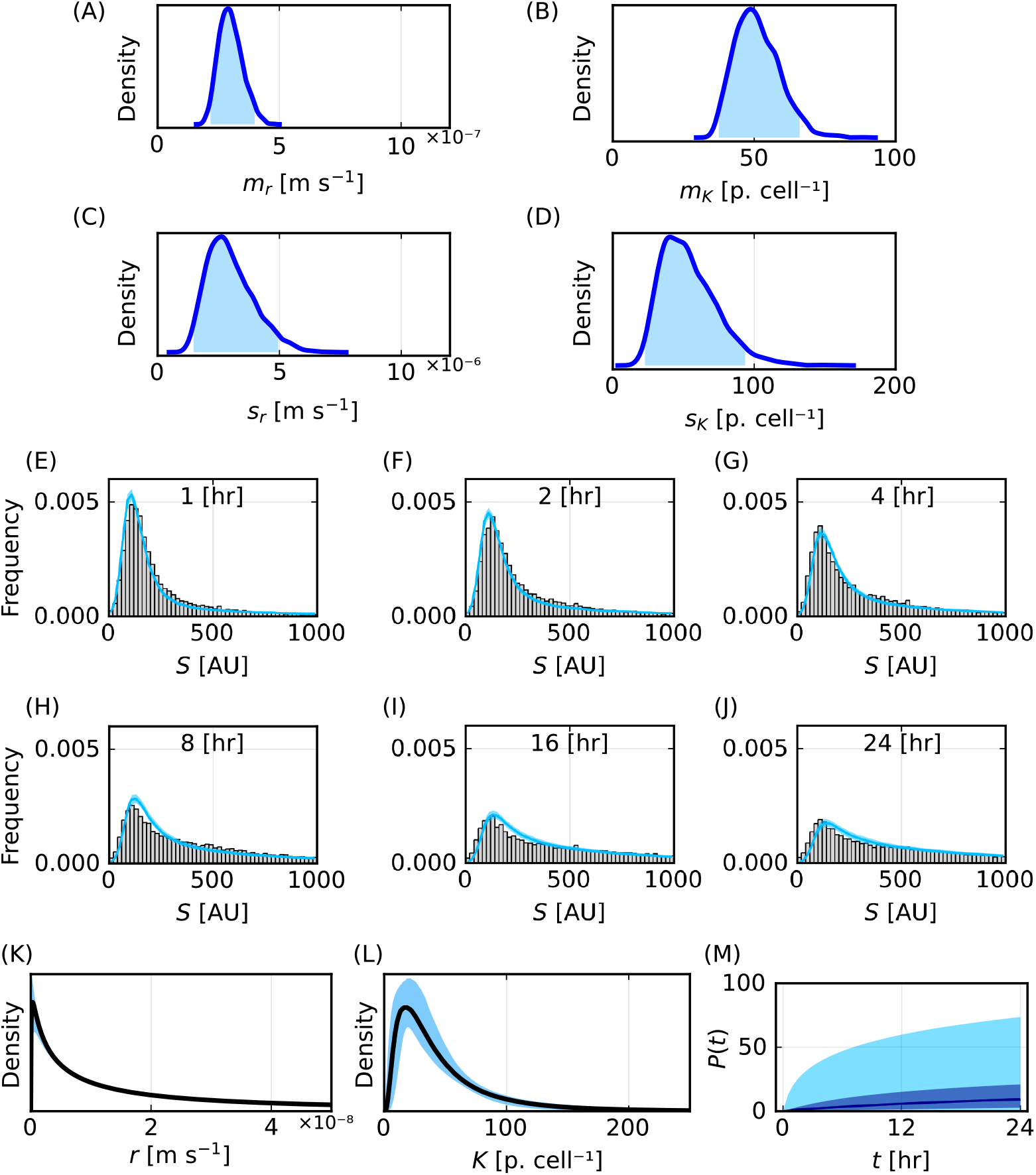
Methods apply to experimental data and reveal uncertainty in predictions of key quantities. (A-D) Posterior distributions for the statistical hyperparameters *m*_*r*_ [m s^*−*1^], *m*_*K*_ [particles cell^*−*1^], *s*_*r*_ [m s^*−*1^], and *s*_*K*_ [particles cell^*−*1^]. The 95% highest posterior density intervals are (2.17 × 10^*−*7^, 3.99 × 10^*−*7^) for *m*_*r*_, (1.48 × 10^*−*6^, 4.95 × 10^*−*6^) for *s*_*r*_, (37.5, 66.1) for *m*_*K*_, and (22.7, 93.6) for *s*_*K*_. (E-J) Histograms of synthetic fluorescence data (grey) and 95% prediction interval of heights of histogram bars from the mathematical model (cyan) for *t* = 1, 2, 4, 8, 16, 24 [hours]. (K)-(L) Inferred distribution for (K) *r* [m s^*−*1^] and (L) *K* [particles cell^*−*1^]. (M) Prediction for *P* (*t*) [particles cell^*−*1^]. In (K)-(M) 95% prediction intervals shaded in blue.

Propagating forward the uncertainty captured by the posterior distributions for the statistical hyperparameters, we observe that posterior predictions for the distribution of fluorescence measurements demonstrate reasonable agreement with the corresponding experimental data (Fig. 7(E-J)). However, the prediction intervals often do not capture the experimental data. We attribute these differences to model misspecification, as we have earlier verified that our mathematical and statistical modelling tools can demonstrate excellent agreement with this type of data and recover known parameters. This model misspecification may arise from the form of the homogeneous mathematical model, the use of lognormal distributions to characterise heterogeneity in *r* and *K*, or the assumptions of additive noise due to cell autofluorescence and multiplicative noise due to particle fluorescence.

We next repeat this analysis for 214 nm PMA-capsules incubated with THP1 cells and 633 nm PMA core-shell particles incubated with THP1 cells (Supp. S2.1). We estimate posterior distributions for the statistical hyperparameters, predictions for the distribution of fluorescence measurements, inferred distributions for *r* and *K*, and predictions for *P* (*t*). We choose the ABC error threshold for the 214 nm and 633 nm data with the Anderson-Darling ABC distance to be 2167 and 1295, which are both higher than the corresponding ABC error threshold for the 150 nm data, which is set to be 847. We make these choices so that the ABC-SMC algorithms terminate in a reasonable timeframe (24 hours). This need for an increased ABC error threshold suggests greater model misspecification for these data.

Analysing these three experimental data sets with two other ABC distance functions, based on the Cramer von Mises statistics and Kolmogorov-Smirnov distance, we find similar agreement between posterior predictions for the distribution of the fluorescence measurements and the corresponding experimental data (Supp. S2.1). Posterior distributions for the statistical hyperparameters and inferred distributions for *r* and *K* obtained using the different ABC distance functions are also similar. This corresponds to predictions for the median of *P* (*t*) demonstrating excellent agreement across the ABC distance functions. However, differences in the posterior distributions result in varied predictions for the upper tails of *P* (*t*). These results suggest that care should be taken when choosing the ABC distance function to interpret these data. For example, if the upper tails of the distribution of *P* (*t*) are important we advise that one should compare results using multiple ABC distance functions.

### 4.5. Comparison to previous methods that focus on point estimates and overlook heterogeneity

Various methods have been employed to infer the dynamics of particle-cell interactions from flow cytometry data [10, 26]. We now compare our new method to recent approaches that seek to quantify the particle-cell association rate, *r*, and carrying capacity, *K*, using mathematical modelling [19, 35, 38, 42]. We focus on these approaches because (i) they assume that the experimental data comprise the same measurements that we have considered thus far: cells incubated with particles at multiple time points, *D*_exp_, cell-only data, *D*_cells_, and particle-only data, *D*_particles_; and (ii) they report results using a useful metric, namely particles per cell [39–41].

Rather than working directly with the entire flow cytometry data, Faria et al. [19] first employ a data transformation to obtain a point estimate of the number of particles per cell at each time point,

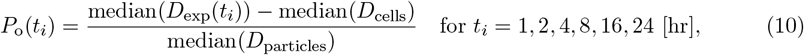

where the subscript ‘o’ denotes observed and *P*_o_(0) = 0. These *P*_o_(*t*_*i*_) are based on the median fluorescence intensity of each flow cytometry data set. While using the median fluorescence intensity to summarise flow cytometry data is a common approach [19, 41], vast amounts of information regarding the heterogeneity of the cell and particle populations are neglected when working solely with median values. For example, for each time point 20,000 cell-level fluorescent measurements in *D*_exp_(*t*_*i*_) are reduced to a single median fluorescence intensity value.

Faria et al. [19] connect these *P*_o_(*t*_*i*_) to the homogeneous mathematical model characterised by two parameters *r* and *K* (Sec. 3.1.1) using the method of least squares (Supp. S1.5). This approach implicitly assumes that measurement/observation errors are additive and follow a Gaussian distribution. It is not clear that these assumptions hold for these data. Furthermore, using the method of least squares, one can only obtain a point estimate for *r* and *K* to characterise the particle-cell interactions. This contrasts our new approach with the heterogeneous mathematical model and ABC methods where we estimate statistical hyperparameters that characterise distributions for *r* and *K* and then generate inferred distributions for *r* and *K*.

For both synthetic data (introduced in Fig. 6) and experimental data (introduced in Fig. 7), point estimates for *r* and *K* that characterise the homogeneous mathematical model obtained by the method of least squares are the same order of magnitude as the modes of the inferred distributions for *r* and *K* generated using the heterogeneous mathematical model (Fig. 8(A,B,D,E,G, H,J,K)). However, while these point estimates for *r* and *K* are the same order of magnitude they do not provide robust accurate estimates of modes of the inferred distributions. Further, these point estimates do not provide robust accurate estimates of the mode of the known distributions for synthetic data (Fig. 8(A,B,D,E,G,H)). We will show that this contributes to poor predictions of the number of particles per cell.

**Figure 8:**
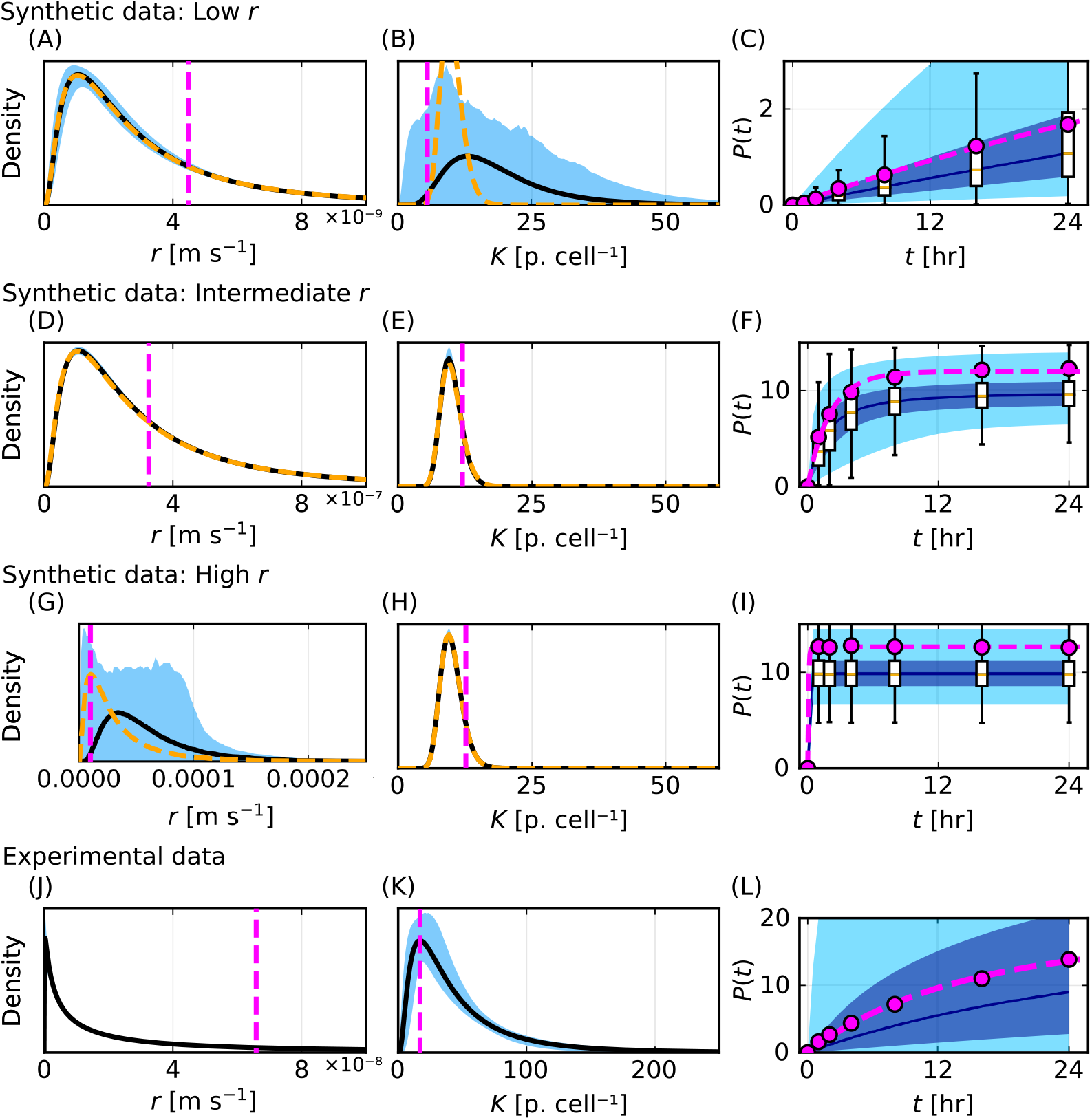
Comparison of point estimate method with inferred distributions and predictions obtained using our new method. (A,D,G,J) Best-fit parameter values for *r* from the homogeneous model with the method of least squares (magenta vertical dashed) compared to inferred distribution for *r* from our new method. (B,E,H,K) Best-fit parameter values for *K* from the homogeneous model with the method of least squares (magenta vertical dashed) compared to inferred distribution for *r* from our new method. (C,F,I,L) *P*_o_(*t*_*i*_) from Eq. (10) (magenta circles) and solution of the homogeneous cell population mathematical model evaluated at the best-fit parameter values for *r* and *K* from the method of least squares (magenta dashed) compared to prediction of *P* (*t*) from our new method.

Simulating the homogeneous mathematical model with the best-fit values of *r* and *K* obtained using the method of least squares, we observe excellent agreement to the transformed data *P*_o_(*t*_*i*_) (from Eq. (10)) for synthetic and experimental data (Fig. 8(C,F,I,L)). However, we find that these predictions from the least squares method systematically overestimate the median of known values of *P* (*t*) from synthetic data. This overestimation appears to arise because control data, *D*_cells_ and *D*_particles_, are right-skewed (Supp. S2.3). Further, for intermediate *r*, these predictions exceed the third quartile of known values of *P* (*t*) from synthetic data at later times (Fig. 8(F)). For high *r*, these predictions exceed the third quartile of the known data at all times (Fig. 8(I)). In contrast, predictions of *P* (*t*) from our new method accurately capture the distribution of *P* (*t*) from synthetic data. These results suggest that our new method is better suited to estimating and predicting heterogeneity and uncertainty in *r, K*, and *P* (*t*).

Previous studies have also recognised that flow cytometry data contains more information than that given by the median fluorescence intensity [32, 35, 47–50, 53]. We now compare our new method to techniques employed in the particle-cell interaction study by Johnston et al. [35]. By generalising Eq. (10) to allow for heterogeneity in *D*_exp_, Johnston et al. estimate the number of particles associated to each cell *j* through time,

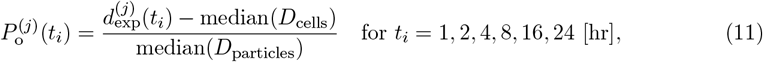

where 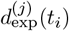 represents the *j*^th^ cell in *D*_exp_ (*t*_*i*_) and we let 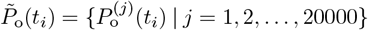. Johnston et al. characterise 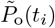at each time point using the mean and standard deviation. While this approach provides two pieces of information to summarise each time point, rather than the one data point provided by Eq. (10), the approach still does not fully exploit the r-488 mation inherent in these data. Furthermore, the approach does not exploit the erogeneity in 489 the cell-only and particle-only control data. Johnston et al. use iterative techniques to fit to the means and standard deviations of 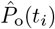 at each time point. This approach provides estimates 491 of lognormal distributions for *r* and *K* that are assumed to characterise these data within a 492 voxel-based mathematical model. Eq. (11) has also been used pre-process experimental data 493 for analysis in [38, 42].

We also note that Eq. (11) can generate non-physical estimates for the number of particles per cell. In particular, when *D*_exp_(*t*_*i*_) is similar to *D*_cells_ = *D*_exp_(0), Eq. (11) can result in many non-physical negative values of 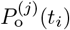. While these non-physical negative values can be discarded, neglecting the negative values can lead to overestimating the number of particles per cell and subsequently lead to inaccurate estimates and predictions. We do not repeat this approach here. Our new method works directly with all data experimental fluorescence measurements and does not require data cleaning or pre-processing procedures.

### 4.6. Identifying optimal experimental time points

Many experimental design choices can influence results and their interpretation. Poor experimental designs may result in poor and inconsistent estimates of particle performance, with the efficacy of some particles being overestimated and the efficacy of others being underestimated. Overestimating particle performance can lead to disappointing results in more complicated experiments and wasted effort and expense. Underestimating particle performance may result in potent particles being overlooked in favour of inferior particles. In contrast, well-designed particle-cell interaction experiments facilitate an improved understanding of the dynamics of particle-cell interactions and support the identification of particles with preferred properties.

We now examine one of the simplest yet most important design choices that can be controlled in these experiments, namely the time points when measurements are obtained. Before an experiment, it is not clear when measurements should be taken to maximise information about particle-cell interactions. To explore how the choice of time points influences results we systematically examine 3003 designs using our heterogeneous mathematical model and established optimal Bayesian design methods for models with intractable likelihoods. For further details see Sec. 3.3.

We examine each of these 3003 experimental designs for three particle-cell scenarios, namely low, intermediate, and high *r*, which are introduced in Fig. 6. We later discuss an overall optimal design for an unknown *r* using data from all three particle-cell scenarios. For each particle-cell scenario, we report the performance of a particular design using a normalised mean utility based on an analysis of twenty synthetic data sets. This mean utility rewards precise estimates of the statistical hyperparameters and is normalised such that the maximum mean utility corresponds to one for each scenario. Throughout this discussion, we highlight nine designs (Fig. 9(A), Supp. Table S4): four simple and naive designs for illustrative purposes (*early, middle, late, uniform*); the time points that we consider in the previous analysis in this study which are those which have been used to collect experimental data previously (*exp r*); an optimal design for each of the three particle-cell scenarios (*optimal low r, optimal intermediate r, optimal high r*); and an overall optimal design (*optimal unknown r*).

**Figure 9:**
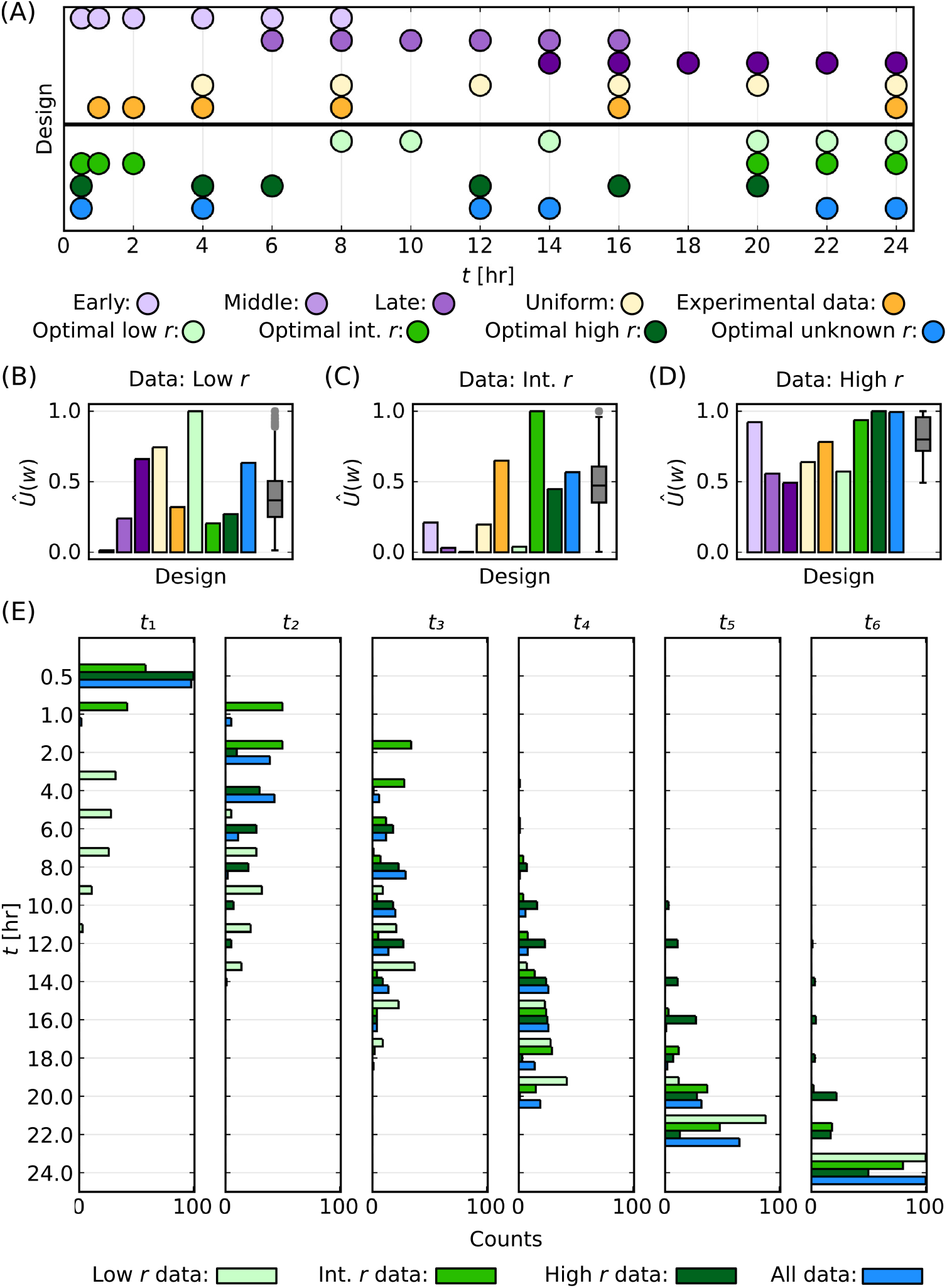
Comparison of 3003 experimental designs across three particle-cell scenarios. (A) Schematic for selected designs. (B-D) Normalised mean utility for selected designs (bars) with a summary for all designs (box plot). Results correspond to data generated for particle-cell scenarios with (B) low *r*, (C) intermediate *r*, and (D) high *r*. (E) Distribution of time points for the top 100 designs for each of the three particle-cell scenarios and for the top 100 designs based on average rank across all three experimental designs. The *i*^th^ time point is denoted *t*_*i*_.

In Fig. 9(B) we present the normalised mean utility function for a scenario with low *r*. We observe that the *optimal low r* design is *t* = 8, 10, 14, 20, 22, 24 [hr]. In Fig. 9(E), we present the distribution of time points for the top 100 designs for low *r*, which demonstrates that later time points correspond to higher utilities for these data. These results agree with expectations. For low *r*, we expect to obtain similar information for *r* across any choice of time points because *P* (*t*) increases approximately linearly throughout the experiment, but we expect to obtain more information about *K* at later times as *P* (*t*) is increasing throughout the experiment. This is consistent with the observation that the *early* design is particularly poor (ranked 3003 out of the 3003 designs) and with the observation that the *exp r* design, which incorporates multiple early time points, is also sub-optimal (ranked 1808 out of the 3003 designs). Further details on the normalised utilities and rankings for the nine selected designs for all scenarios are presented in Supp. Table S4.

The *optimal intermediate r* design comprises multiple measurements at early times and multiple measurements at late times (*t* = 0.5, 1, 2, 20, 22, 24 [hr]) (Fig. 9(A)). The top 100 designs for intermediate *r* follow a similar structure (Fig. 9(E)). The structure of these top designs is expected since we seek to capture the prominent roles of *r* at early time and *K* at late time.

The *optimal high r* design corresponds to measurements throughout the experiment (*t* = 0.5, 4, 6, 12, 16, 20 [hr]) (Fig. 9(A)). The top 100 designs for high *r* follow a similar structure (Fig. 9(E)). We expect this structure for these top designs since data for high *r* is consistent across all time points. Many designs result in a similar normalised mean utility with the lowest normalised mean utility for high *r* equal to 0.492 whereas the lowest normalised mean utility for low and intermediate *r* are 0.014 and 0.003, respectively (Fig. 9(B-D)).

All optimal designs maximise the normalised mean utility for their respective scenarios. Furthermore, the *optimal low r* and *optimal int r* designs correspond to outliers in the 3003 designs with respect to the normalised mean utility (Fig. 9(B-D)). However, a design that is optimal in one scenario is not necessarily optimal for a different scenario. For example, the *optimal low r* design is a poor design for intermediate *r* (ranked 2922 out of 3003 designs) and high *r* (ranked 2914 out of 3003 designs).

Typically particle-cell interaction experiments are performed to infer the particle-cell association rate, so we often do not know whether the experiment corresponds to a scenario with low, intermediate, or high *r*. Therefore, we compute an overall optimal design for an unknown *r* based on an average of the ranking of designs across the three particle-cell scenarios. This *optimal unknown r* design has the highest average ranking, which is 418, and comprises measurements throughout the experiment (*t* = 0.5, 4, 12, 14, 22, 24 [hr]) (Fig. 9(A)). This design corresponds to rankings of 298, 939, and 16 for data with low, intermediate, and high *r*, respectively. These rankings exceed 90%, 69%, and 99% of the 3003 designs for low, intermediate, and high *r*, respectively (Fig. 9(B-D)).

The overall top 100 designs for unknown *r* across all data follow a similar structure to the *optimal unknown r* design (Fig. 9(E)). The *optimal low r, optimal intermediate r*, and *optimal high r* designs are not found in the overall top 100 designs at positions 2430, 855, and 1000, respectively. The *exp r* design is positioned 1430 in the overall ranking, with an average ranking of 1438 due to rankings of 1808, 565, and 1942 for data with low, intermediate, and high *r*, respectively. These results suggest that while the design that has been previously used to collect data is in the top 20% of designs for scenarios with intermediate *r*, there are 1429 designs better suited to explore scenarios when *r* is unknown.

## 5. Conclusion

In this study, we use mechanistic mathematical modelling, statistical modelling, and techniques from optimal Bayesian design to examine routinely collected flow cytometry measurements from nano-engineered particle-cell interaction experiments. We exploit previously overlooked and heterogeneity in routine measurements that form the particle-only control data, cell-only control data, and time course flow cytometry data using a novel heterogeneous ordinary differential equation-based mathematical model. This approach allows us to reveal and quantify heterogeneity and associated uncertainty in key biological parameters that characterise particle-cell interaction experiments. The approach also allows us to generate predictions of key quantities that are challenging to observe by experimentation alone, including the time evolution of the number of particles per cell. While many studies quantify the particle-cell interactions by assessing the percentage of cells positive for the presence of particles, using our tools we can generalise this metric to estimate how many particles are present in each cell. Obtaining such insights at scale with the latest experimental technology is challenging. Previous methods that assume homogeneous cell populations are also unable to provide such insights.

We apply and verify that our new tools perform as desired in various particle-cell scenarios. Classifying these scenarios using flow cytometry data without modelling is challenging. Instead, we identify different experimental regimes using predictions of the time evolution of the number of particles per cell from our mathematical modelling approach. To identify model parameters we seek a regime where we obtain multiple measurements of the initial increase in the number of particles per cell and multiple measurements of the number of particles per cell close to carrying capacity. Scenarios that do not include either of these types of measurements result in increased uncertainty and lack of identifiability for the particle-cell association rate and carrying capacity-type parameter, respectively.

Applying our tools to experimental data with particles that range from 150 nm to 633 nm incubated with a human leukaemia monocytic cell line, we estimate key biological parameters and make predictions that capture the heterogeneity in the data. We demonstrate that these methods improve on previous methods that assume homogeneous cell populations and those that rely on pre-processing calculations that can produce non-physical quantities. These previous methods also use the method of least squares and iterative techniques to obtain point estimates of key parameters that do not capture the heterogeneity in the data. While the uncertainty in these point estimates could be estimated and propagated forward to generate predictions [65], it is not clear what the most appropriate measurement/observation error model should be. The approach that we employ, with a heterogeneous mathematical model and approximate Bayesian computation, circumvents this challenge and, among many advantages, explicitly captures heterogeneity.

Model predictions for the flow cytometry data demonstrate reasonable agreement with the corresponding experimentally measured data for multiple ABC distance functions. In particular, predictions for the number of particles per cell from different ABC distance functions closely agree about the median. As we verify that our methods recovers known quantities for synthetic data, we attribute differences to model misspecification [62, 63], which is near impossible to avoid in such a complex system as particle-cell interactions. To explore model misspecification, one could revisit the assumptions of the mathematical models and develop more sophisticated models, likely with more parameters, but this may lead to further parameter identifiability challenges. One could also revisit the assumption that we capture all measurement noise in the flow cytometry time course data by sampling from the experimental control data, and allow for additional measurement noise by extending our ABC method to an exact ABC inference technique [64]. In the first instance, one can start to observe the impact of model misspecification by generating results using multiple ABC distance functions. This may be particularly insightful when seeking estimates of the statistical hyperparameters and predictions for the upper tails of the distribution for the number of particles per cell.

Using optimal experimental design techniques with our novel model, we identify optimal time points for multiple particle-cell scenarios. This analysis systematically examines 3003 designs and rewards designs that obtain precise estimates of the parameters that characterise particle-cell interactions. We focus on time points as they are simple to vary experimentally and vital for accurate parameter estimates. We choose the set of possible time points based on previous data and practical considerations regarding the frequency of measurements with manual experimental procedures. Automated experimental procedures would facilitate more frequent measurements. When early time measurements do not capture the initial increase in particles per cell, increasing the temporal resolution at early times would facilitate a transition from a regime of partial identifiability to a regime where parameters are identifiable. When late time measurements of the number of particles per cell are not close to carrying capacity, extending the experimental duration would also facilitate a transition to a regime where model parameters are identifiable. This would, however, introduce additional complications; for example, cell proliferation and media changes are expected to be more influential. The methodology that we present here is well-suited to incorporate such additional mechanisms.

Overall, our results suggest that it is essential to consider heterogeneity even when interpreting the simplest of controlled *in vitro* nano-engineered particle-cell interaction experiments. We quantify this heterogeneity in the particle-cell association rate and carrying capacity, but we do not claim to know the mechanisms that drive this heterogeneity. Exploring the mechanisms that give rise to such heterogeneity would be interesting. The methodology we present in this study is well-suited to analyse other combinations of particles and cell types in well-mixed media, such as with suspension and adherent cell lines subject to stirring. The methodology is suitable for extensions where spatial gradients of particles form in the media, such as for adherent cells in an unstirred media, and this would involve extending the mathematical model to a partial differential equation-based model that incorporates the role of particle transport [19, 35]. We also contribute to the growing literature demonstrating the advantages of working with all flow cytometry data rather than summary statistics, such as the median fluorescence intensity [47–50, 53]. We show that experimental data can exhibit dramatic cell-cell variability in particle-cell association rates, carrying capacity, and associated particles, and the influence of these findings on target applications, such as medical diagnostics, biomedical imaging, and targeted drug delivery, is worthy of further investigation.

## Data and code availability

We analyse previously published data from [19] available on FigShare. This includes fluorescence data from Flow Cytometry Standard (FCS) files and experimental details from INI files. Raw FCS data files were converted to CSV using https://floreada.io/. Key computer code implemented in Julia is available on GitHub (https://github.com/ryanmurphy42/Murphy2025NanoUQ).

## Author’s contributions

All authors conceived and designed the study. RJM performed the research and drafted the article. All authors provided comments and approved the final version of the manuscript.

## Acknowledgements

This research was supported by The University of Melbourne’s Research Computing Services and the Petascale Campus Initiative. We thank Dr David J. Warne and Dr Alexander P. Browning for helpful discussions.

