## Supplementary Material for "Quantifying biological heterogeneity in nano-engineered particle-cell interaction experiments"

**S1 Methods** **3**

### S1 Methods

#### S1.1 Additional experimental details

The experimental data, $\hat{D}_{\text{exp}}$, $\hat{D}_{\text{cells}}$, and $\hat{D}_{\text{particles}}$, described in Sec. 2, are collected using different photomultiplier voltage settings (500, 500, and 600 [V], respectively) [1]. To account for the different voltage settings, we calibrate all data to the same reference voltage, $V_{\text{ref}}$ [1]. We set the reference voltage equal to the voltage used to obtain the particle-only measurements, i.e. $V_{\text{ref}} = V_{\text{particles}}$, and set $D_{\text{particles}} = \hat{D}_{\text{particles}}$. For the cell-only control data and time-course data we set $D_{\text{cells}} = C_{\text{pmt}}(V_{\text{cells}}, V_{\text{ref}})\hat{D}_{\text{cels}}$ and $D_{\text{exp}} = C_{\text{pmt}}(V_{\text{exp}}, V_{\text{ref}})\hat{D}_{\text{exp}}$, respectively, where $C_{\text{pmt}}(V, V_{\text{ref}}) = \exp(0.016(V_{\text{ref}} - V))$ is a previously calibrated correction factor. All flow cytometry measurements are recorded using the 638 (Red)Peak detected signal [1].

#### S1.2 Measured and fixed parameters

The homogeneous and heterogeneous mathematical models are characterised by four parameters, $(C, S, V, u_0)$, that are directly measured in the experiments. Therefore, we assume that these parameters are fixed constants. Here, we present the formulae and values for these parameters. Formulae are written in terms of variables that are included in the INI files that summarise the experiments and are available via [1] (`total_cells` $= 100,000$, `surface_area_per_cell_in_m2` $= 530 \times 10^{-12}$, `height_in_m` $= 0.00262$, `width_in_m` $= 0.0221$, `particles_per_cell` $= 100$). Throughout, we use the fact that we consider a suspension cell line incubated with particles, which means that the entire surface of each cell is exposed to the media that contains particles.

| Variable | Units | Description | Formula (from INI file variables) | Value |
|---|---|---|---|---|
| $C$ | - | Fractional surface coverage of cells | 1 | 1 |
| $S$ | $m^2$ | Cell boundary surface area | `total_cells` $\times$ `surface_area_per_cell_in_m2` | $5.31 \times 10^{-5}$ |
| $V$ | $m^3$ | Media volume | $\pi \times$ `height_in_m` $\times$ `width_in_m`$^2/4$ | $1.01 \times 10^{-6}$ |
| $u_0$ | $m^{-3}$ | Initial number density of particles per cell | `particles_per_cell`$/V$ | $9.95 \times 10^{7}$ |

**Table S1: Measured and fixed parameters, their units, description, formula, and values.**

#### S1.3 Heterogeneous mathematical model

In the main manuscript, we present the approximate solution to the heterogeneous mathematical model that we use for parameter estimation, practical identifiability analysis, and prediction. Here, we present the system of ordinary differential equations that defines the heterogeneous mathematical model and demonstrate that the computationally efficient approximate solution demonstrates excellent agreement with the solution to the heterogeneous model for experimentally relevant parameter regimes.

##### S1.3.1 Derivation of the heterogeneous mathematical model

Previous studies have demonstrated that the particle-cell association rate, $r$, and carrying capacity, $K$, exhibit significant variability between cells subject to the same experimental conditions [2]. We now make this same modelling assumption and generalise the previously published homogeneous model described in Sec. 3.1.1 of the main manuscript and [1]. In particular, we make a common assumption for non-negative biological parameters and assume that $r$ and $K$ are lognormally distributed [3–5],

$$\begin{aligned} r &\sim \text{LogNormal}(m_r, s_r), \\ K &\sim \text{LogNormal}(m_K, s_K). \end{aligned} \tag{S.1}$$

As we are allowing for cell-cell heterogeneity in $r$ and $K$, in the following we consider the total number of particles in the well-mixed media at time $t$, denoted $U(t)$ [particles cell$^{-1}$] with initial condition $U(0) = U_0$ [particles cell$^{-1}$].

We let $N$ denote the total number of cells in the well and $\tilde{N}$ denote the number of cells that we will simulate. We choose $\tilde{N} = 20,000$ to equal the number of experimental measurements per time point and assume that this is sufficiently large to numerically approximate the $r$-$K$ distribution. Assuming that the dynamics for particle-cell interactions are the same as in the homogeneous model, the total number of particles associated with cell $j$ for $j = 1, 2, \ldots, \tilde{N}$, denoted $P^{(j)}(t)$, is governed by,

$$\frac{\mathrm{d}P^{(j)}(t)}{\mathrm{d}t} = \frac{\tilde{C}Sr_j}{V}\left(\frac{K_j - P^{(j)}(t)}{K_j}\right)U(t), \tag{S.2}$$

where $r_j$ and $K_j$ denote the particle-cell association rate and carrying capacity of cell $j$, respectively, and the scaled fractional surface coverage of cells is

$$\tilde{C} = C/N. \tag{S.3}$$

Analogous to Eq. (2), by conservation of the total number of particles and and assuming that there is no particle degradation during the experiment, the total number of associated particles at time $t$ is equal to the difference between the initial number of particles in the media and the total number of particles in the media at time $t$,

$$\frac{N}{\tilde{N}}\sum_{j=1}^{\tilde{N}} P^{(j)}(t) = U_0 - U(t), \tag{S.4}$$

where we assume that each of the $\tilde{N}$ simulated cells represents $N/\tilde{N}$ of the total number of cells in the experimental well.

Differentiating Eq. (S.4) with respect to time and substituting in Eq. (S.2) and Eq. (S.3), we obtain an equation for the time evolution of $U(t)$,

$$\frac{\mathrm{d}U(t)}{\mathrm{d}t} = -\frac{N}{\tilde{N}} \sum_{j=1}^{\tilde{N}} \frac{\mathrm{d}P^{(j)}(t)}{\mathrm{d}t} = -\frac{1}{\tilde{N}} \sum_{j=1}^{\tilde{N}} \frac{CSr_j}{V} \left( \frac{K_j - P^{(j)}(t)}{K_j} \right) U(t). \tag{S.5}$$

Eqs. (S.2) and (S.5) form a system of $\tilde{N} + 1$ ordinary differential equations for $U(t)$ and $P^{(j)}(t)$ for $j = 1, 2, \dots \tilde{N}$. Solving this system of equations can be computationally expensive for $\tilde{N} = 20,000$ and renders inference challenging. To address this challenge, we determine an approximate solution to this heterogeneous model that can be obtained significantly faster and supports efficient inference.

Before proceeding to the approximate solution and to enable direct comparisons to the homogeneous model, we convert the equations given in terms of the total number of particles to equations in terms of the concentration of particles per cell $u(t)$ in the media [particles cell$^{-1}$ m$^{-3}$],

$$u(t) = \frac{U(t)}{NV}. \tag{S.6}$$

Eqs. (S.2) and (S.5), written in terms of $u(t)$, are then

$$\frac{\mathrm{d}P^{(j)}(t)}{\mathrm{d}t} = CSr_j \left( \frac{K_j - P^{(j)}(t)}{K_j} \right) u(t), \quad \text{for} \quad j = 1, 2, \dots, \tilde{N}, \tag{S.7}$$

and

$$\frac{\mathrm{d}u(t)}{\mathrm{d}t} = -\frac{1}{\tilde{N}} \sum_{j=1}^{\tilde{N}} \frac{CSr_j}{V} \left( \frac{K_j - P^{(j)}(t)}{K_j} \right) u(t). \tag{S.8}$$

In Sec. S1.3.4 we numerically solve Eqs. (S.7) and (S.8) to verify the accuracy of the approximate solution to the heterogeneous model that is described in Sec. S1.3.2). Specifically, we numerically solve Eqs. (S.7) and (S.8) using the DifferentialEquations.jl and Sundials.jl Julia packages [6–8] and the CVode Backward Differentiation Formula with a GMRES linear solver, absolute error tolerance equal to $1 \times 10^{-10}$, and relative error tolerance equal to $1 \times 10^{-10}$.

##### S1.3.2 Approximate solution to the heterogeneous model

In experiments, the total number of particles in the well-mixed media is simple to vary experimentally and typically chosen sufficiently large such that many particles will remain in the media throughout the experiment. Therefore, one expects that $u(t) \approx u_0$ throughout the duration of the experiment. For improved accuracy and to approximate the influence of cell-cell heterogeneity, we approximate $u(t)$ with

$$\bar{u}(t) = \frac{1}{M} \sum_{m=1}^{M} u_m(t), \tag{S.9}$$

where $u_m(t)$ is the concentration of particles per cell in the well-mixed media obtained by evaluating the analytical solution of the homogeneous model (Eq. (3)) at time $t$ with parameters $r_m$ and $K_m$ sampled from the probability distributions defined in Eq. (S.1). We set $M = 100$ throughout based on pilot simulations exploring a tradeoff between accuracy and computational efficiency.

Given $\bar{u}(t)$, Eqs. (S.7)-(S.8) are decoupled with solution,

$$P^{(j)}(t) = K_j \left(1 - \exp\left(-\frac{CSr_j}{K_j}\int_0^t \bar{u}(\hat{t})\ \mathrm{d}\hat{t}\right)\right), \quad \text{for} \quad j = 1, 2, \ldots, \tilde{N}, \tag{S.10}$$

where we estimate the integral efficiently and numerically using the trapezoid rule with node spacing equal to 0.5 hours.

##### S1.3.3 Reduction of heterogeneous model to homogeneous model

Here, we demonstrate that the heterogeneous model reduces to the previously published homogeneous model under the assumption of identical cells.

Setting $r = r_j$, $K = K_j$, and $P(t) = P^{(j)}(t)$ in Eq. (S.8) gives,

$$\frac{\mathrm{d}u(t)}{\mathrm{d}t} = -\frac{1}{\tilde{N}}\sum_{j=1}^{\tilde{N}} \frac{CSr}{V}\left(\frac{K - P(t)}{K}\right)u(t) = -\frac{CSr}{V}\left(\frac{K - P(t)}{K}\right)u(t), \tag{S.11}$$

which is identical to Eq. (1) in the homogeneous model. Similarly, setting $r = r_j$, $K = K_j$, and $P(t) = P^{(j)}(t)$ in Eqs. (S.7) gives

$$\frac{\mathrm{d}P(t)}{\mathrm{d}t} = CSr\left(\frac{K - P(t)}{K}\right)u(t), \tag{S.12}$$

which is identical to the equation for $P(t)$ in the homogeneous model if one differentiates Eq. (2) with respect to time and applies Eq. (1).

| Variable/Parameter | Units | Description | Mathematical model |
|---|---|---|---|
| $C$ | - | Fractional surface coverage of cells | Homogeneous/Heterogeneous |
| $\tilde{C}$ | - | Scaled fractional surface coverage of cells | Heterogeneous |
| $K$ | particles cell$^{-1}$ | Cell carrying capacity for particles | Homogeneous |
| $K_j$, $K_m$ | particles cell$^{-1}$ | Cell $j$,$m$ carrying capacity for particles | Heterogeneous |
| $M$ | - | Number of simulations to approximate $\bar{u}(t)$ | Heterogeneous |
| $m_r$ | m s$^{-1}$ | Mean particle-cell association rate | Heterogeneous |
| $m_K$ | particles cell$^{-1}$ | Mean cell carrying capacity for particles | Heterogeneous |
| $N$ | - | Number of cells in the experimental well | Heterogeneous |
| $\tilde{N}$ | - | Number of simulated cells in the experimental well | Heterogeneous |
| $P(t)$ | particles cell$^{-1}$ | Number of associated particles per cell at time $t$ | Homogeneous |
| $P^{(j)}(t)$ | particles cell$^{-1}$ | Number of associated particles per cell at time $t$ for simulated cell $j$ | Heterogeneous |
| $r$ | m s$^{-1}$ | Particle-cell association rate | Homogeneous |
| $r_j$, $r_m$ | m s$^{-1}$ | Particle-cell association rate for cell $j$, $m$ | Heterogeneous |
| $S$ | m$^2$ | Cell boundary surface area | Homogeneous/Heterogeneous |
| $s_r$ | m s$^{-1}$ | Standard deviation for particle-cell association rate | Heterogeneous |
| $s_K$ | particles cell$^{-1}$ | Standard deviation for cell carrying capacity for particles | Heterogeneous |
| $t$ | hr | Time | Homogeneous/Heterogeneous |
| $u(t)$ | particles cell$^{-1}$ m$^{-3}$ | Concentration of particles per cell in media | Homogeneous |
| $u_0$ | particles cell$^{-1}$ m$^{-3}$ | Initial concentration of particles per cell in media | Homogeneous |
| $U(t)$ | particles | Total number of particles in media at time $t$ | Heterogeneous |
| $U_0$ | particles | Initial total number of particles in the media | Heterogeneous |
| $\bar{u}(t)$ | particles cell$^{-1}$ m$^{-3}$ | Approximate concentration of particles per cell in media | Heterogeneous |
| $u_m(t)$ | particles cell$^{-1}$ m$^{-3}$ | Concentration of particles per cell in media from homogeneous model | Heterogeneous |
| $V$ | m$^3$ | Media volume | Homogeneous/Heterogeneous |

**Table S2: Parameters/variables, their units, and descriptions.**

##### S1.3.4 Verification of the approximate solution to the heterogeneous model

We now verify that the approximate solution to the heterogeneous model (Eqs. (S.9) and (S.10)) demonstrates excellent agreement with the solution of the heterogeneous model (obtained by solving Eqs. (S.7) and (S.8)) for experimentally relevant parameter regimes.

We first present results for the typical experimental scenario where the total number of particles in the well-mixed media is chosen sufficiently large such that many particles will remain in the media throughout the experiment. In particular, we assume that there are initially 100 particles per cell, define the lognormal distribution for $K$ [particles cell$^{-1}$] with $m_K = 40.0$ and $s_K = 10.0$, and define the lognormal distribution for $r$ [m s$^{-1}$] with $m_r = 4.0 \times 10^{-6}$ and $s_r = 1.0 \times 10^{-6}$. This corresponds to predictions for the number of particles per cell from the heterogeneous model (Eqs. (S.7) and (S.8)) remaining below the initial number of particles per cell (Fig. S1(A)). In this scenario, the approximate solution to the heterogeneous model demonstrates excellent agreement with the corresponding solution to the heterogeneous model for both the number density of particles in the media (Fig. S1(C)) and the number of particles per cell (Fig. S1(E)).

We now present results for a non-typical experimental scenario where most of the particles initially in the media associate with cells. It is important to note that this scenario does not apply to the experimental data that we analyse. The following example is included primarily to demonstrate that it is possible for the approximate solution to the heterogeneous model to demonstrate poor agreement with the corresponding solution to the heterogeneous model. We maintain the assumptions that there are initially 100 particles per cell and that the lognormal distribution for $r$ [m s$^{-1}$] is characterised by $m_r = 4.0 \times 10^{-6}$ and $s_r = 1.0 \times 10^{-6}$, however we vary the lognormal distribution for $K$ [particles cell$^{-1}$] and set $m_K = 110.0$ and $s_K = 50.0$. This results in predictions for the number of particles per cell from the heterogeneous model (Eqs. (S.7) and (S.8)) approaching and exceeding the initial number of particles per cell (Fig. S1(B)). Here, we do not observe good agreement between the approximate solution to the heterogeneous model and the corresponding solution to the heterogeneous model (Fig. S1(D,F)). This poor agreement is because the approximate solution overestimates the number of remaining particles in the media as $t$ increases.

For the experimental and synthetic data that we analyse, the approximate solution to the heterogeneous model demonstrates excellent agreement with the corresponding solution to the heterogeneous model for both the concentration of particles per cell in the media and the number of particles per cell. This is because the total number of particles in the well-mixed media is sufficiently large such that many particles will remain in the media throughout the experiment. As a representative example, we present results for experimental data with the 150 nm PMA core-shell particles, discussed in Sec. 4.4. Examining each sample $k$ in the posterior distribution for the statistical hyperparameters $(m_r^{(k)}, s_r^{(k)}, m_K^{(k)}, s_K^{(k)})$, we compute an average absolute error, $e_k$, across all time points and all $\tilde{N}$ simulated cells,

$$e^{(k)} = \frac{1}{6} \sum_{i=1}^{6} \frac{1}{\tilde{N}} \sum_{j=1}^{\tilde{N}} \left| P^{(j)}\left(t_i; u(t_i), m_r^{(k)}, s_r^{(k)}, m_K^{(k)}, s_K^{(k)}\right) - P^{(j)}\left(t_i; \bar{u}(t_i), m_r^{(k)}, s_r^{(k)}, m_K^{(k)}, s_K^{(k)}\right) \right|. \tag{S.13}$$

This error is small for all samples in the posterior distribution (Fig. S2(A,B)). We also find excellent agreement between the solution from the heterogeneous model and the approximate solution to the heterogeneous model for the parameters values with the maximum error (Fig. S2(C,D)). We find similar excellent agreement for the synthetic data studies and for experimental data with the 214 nm PMA-capsules and 633 nm PMA core-shell particles.

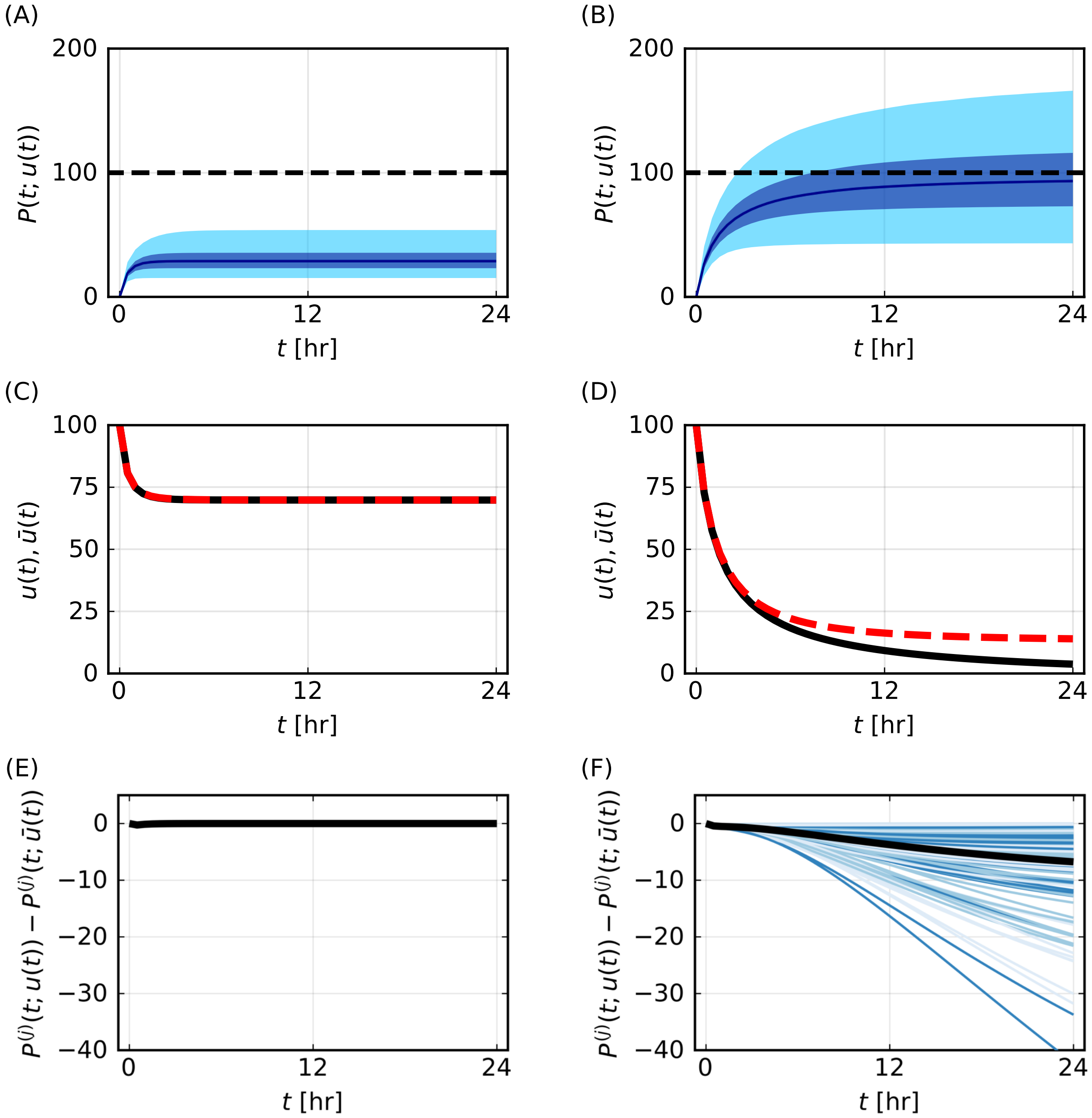

**Figure S1: Exemplar comparison of solutions to the heterogeneous model (Eqs. (S.7) and (S.8)) and the corresponding approximate solutions to the heterogeneous model (S.7) and (S.8)).** (A,B) Predictions for the number of particles per cell, $P(t)$ [particles cell$^{-1}$], from the heterogeneous model. Initial number of particles per cell shown as horizontal black dashed line. (C,D) Comparison of $u(t)$ (black) from the heterogeneous model to $\bar{u}(t)$ (red-dashed) from the approximate solution to the heterogeneous model. (E,F) Comparison of difference between the solution of $P^{(j)}(t; u(t))$ from the heterogeneous model and the corresponding approximate solution of $P^{(j)}(t; \bar{u}(t))$ for $j = 1, 2, \ldots, 200$ randomly sampled simulated cells (multiple shades of blue) with the mean difference (black). Parameter values for (A,C,E) are $(m_r,\ s_r,\ m_K,\ s_K) = (4.0 \times 10^{-6},\ 1.0 \times 10^{-6},\ 40.0,\ 10.0)$ and for (B,D,F) are $(m_r,\ s_r,\ m_K,\ s_K) = (4.0 \times 10^{-6},\ 1.0 \times 10^{-6},\ 110.0,\ 50.0)$.

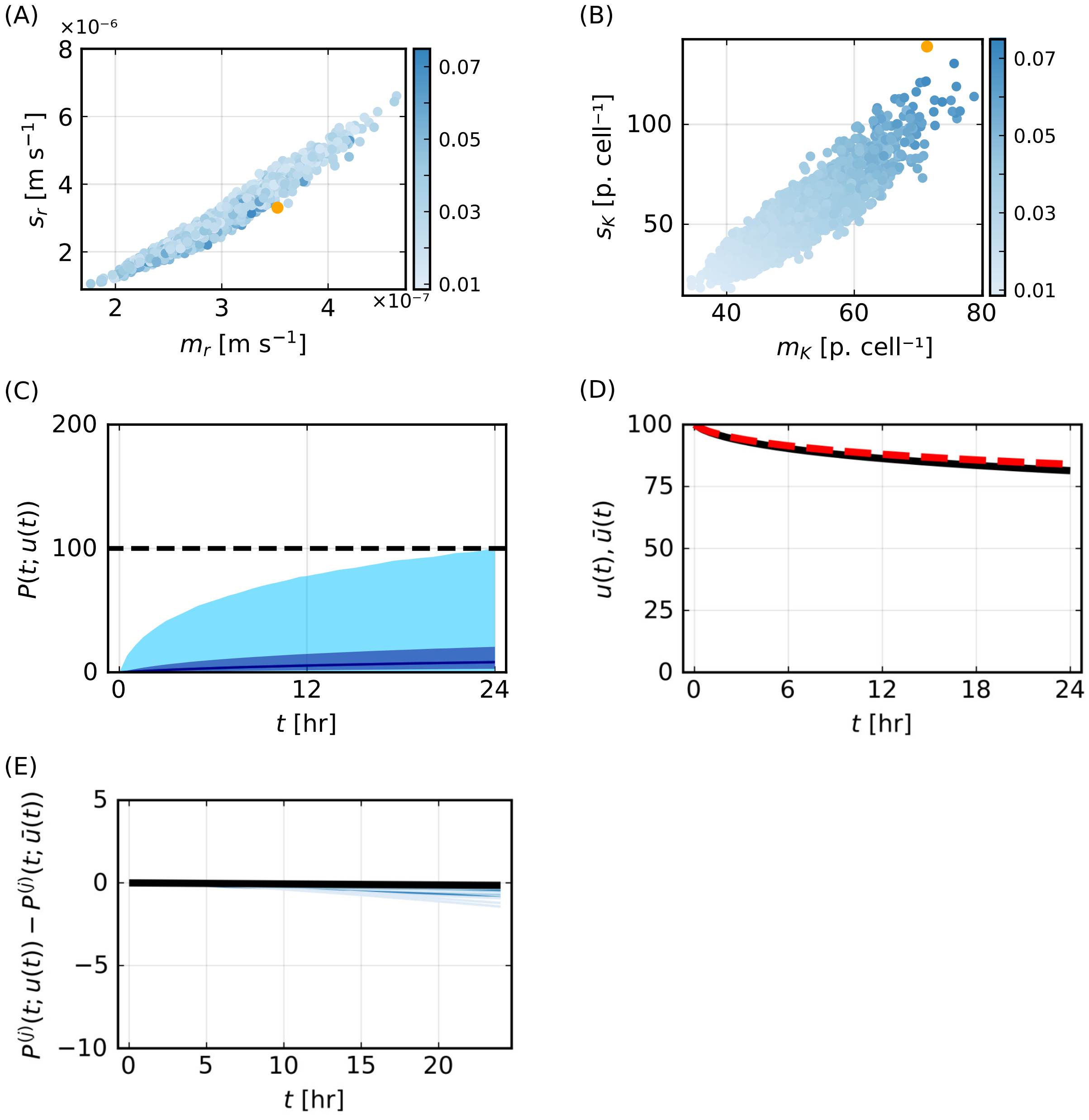

**Figure S2: Comparison of solutions to the heterogeneous model and the corresponding approximate solutions to the heterogeneous model using parameter values from the ABC posterior distribution for the 150 nm PMA core-shell particles experimental data.** (A,B) Average absolute error from Eq. (S.13) for each sample of the posterior distribution for parameters that characterise the distribution of (A) $r$ and (B) $K$. In (A-B) the colour bar represents the error defined in Eq. (S.13) and the orange circle represents the parameter values $(m_r^{(k)}, s_r^{(k)}, m_K^{(k)}, s_K^{(k)}) = (3.5 \times 10^{-7}, 3.3 \times 10^{-6}, 71.4, 139.0)$ with the maximum error, $e^{(k)} = 0.075$. Results in (C-E) use the parameter values with the maximum error. (C) Prediction of number of particles per cell, $P(t)$, simulated using parameter values. Initial number of particles per cell shown as horizontal black dashed line. (D) Comparison of $u(t)$ (black) from the full model to $\bar{u}(t)$ (red-dashed) from the approximate solution. (E) Comparison of difference between the solution of $P^{(j)}(t; u(t))$ from the full model and the corresponding approximate solution of $P^{(j)}(t; \bar{u}(t))$ for $i = 1, 2, \ldots, 200$ (multiple shades of blue) with the mean difference (black).

#### S1.4 Parameter estimation, practical identifiability analysis, and prediction for the heterogeneous mathematical model

In Sec. 3 of the main manuscript we outline the approximate Bayesian computation (ABC) approach that we use for parameter estimation, practical identifiability analysis, and prediction. Here we present further details. In brief, we seek parameter values that minimise the difference between the *experimentally measured* flow cytometry data and the corresponding *simulated* data from the approximate solution to the heterogeneous mathematical model (Fig. S3).

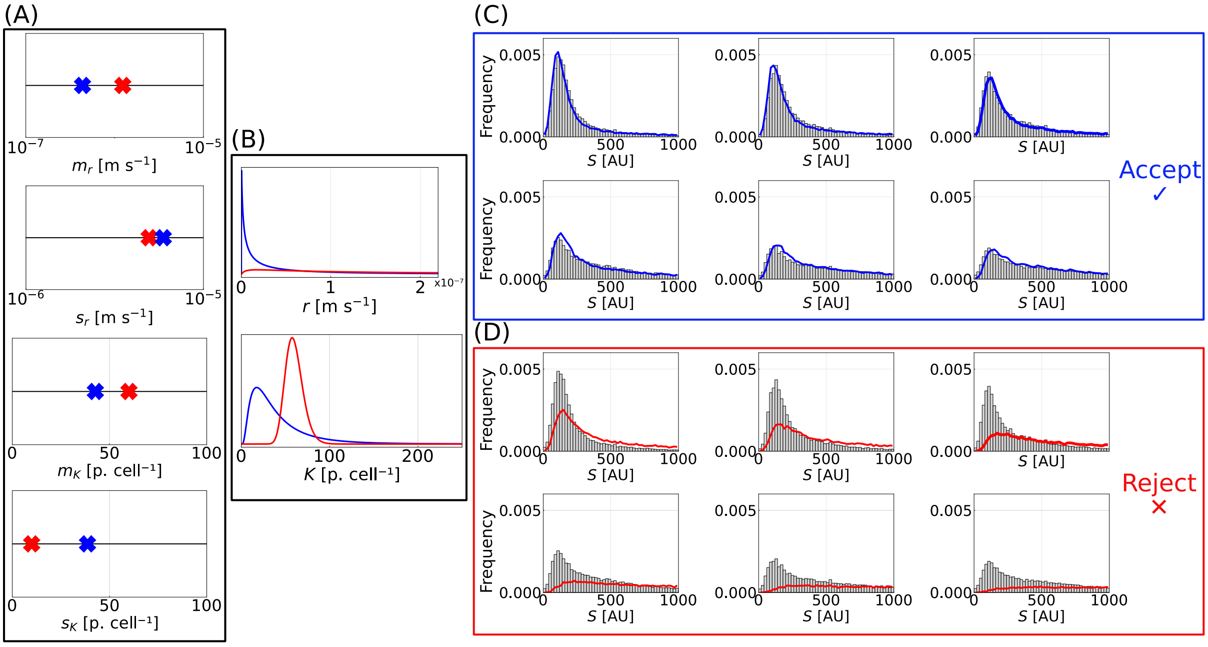

**Figure S3: Schematic of ABC inference methods.** (A) Sample uniform priors for the statistical hyperparameters. Blue/red crosses represent values that result in synthetic data with good/poor agreement to the experimental data. (B) Lognormal distributions for $r$ and $K$ based on the sampled hyperparameter values. (C) Simulate the mathematical model to generate simulated data (blue) and compare to the experimental data (grey histogram). Accept the sampled hyperparameter values if the simulated and experimental data are close and reject otherwise. (D) represents a repeat of (C) with sampled hyperparameters that are rejected.

*Forward simulation of the mathematical model.* Experimental data comprises of flow cytometry measurements $D_{\text{exp}} = \{D_{\text{exp}}(t_i) \mid i = 1, 2, \dots, 6\}$. To simulate synthetic data comparable to $D_{\text{exp}}$ for fixed values of statistical hyperparameters, we use the approximate solution to the heterogeneous mathematical model, cell-only control experimental data, and particle-only control experimental data (Sec. 3).

Fixing values of the statistical hyperparameters (Fig. S3(A)) defines lognormal distributions for $r$ and $K$ (Fig. S3(B)). For each time point $t_i$, for $i = 1, 2, \dots, 6$, we sample these lognormal distributions $\tilde{N} = 20,000$ times. Once the samples of the lognormal distributions for $r$ and $K$ are determined, we simulate the number of particles per cell, $P^{(j)}(t_i) = P^{(j)}(t_i; r^{(j)}, K^{(j)})$ for $j = 1, 2, \dots, \tilde{N}$ and each $t_i$, using the approximate solution to the heterogeneous mathematical model. As explained in Sec. 3.2, we then convert each $P^{(j)}(t_i)$ to a simulated fluorescence value by sampling from the cell-only and particle-only control experimental data. For each time point we denote the simulated data $D_{\text{sim}}(t_i) = \{d_{\text{sim}}^{(j)} \mid j = 1, 2, \dots \tilde{N}\}$. We denote simulated data over all time points as $D_{\text{sim}} = \{D_{\text{sim}}(t_i) \mid \imath = 1, 2, \dots, 6\}$.

*Distance function.* Given measured experimental data, $D_{\text{exp}}$, and simulated data, $D_{\text{sim}}$, we introduce a distance function based on the Anderson-Darling statistic [9, 10] to compare how *close* they are over all time points,

$$\mathcal{D}_{\text{AD}}(D_{\text{exp}}, D_{\text{sim}}) = \sum_{i=1}^{6} \hat{\mathcal{D}}_{\text{AD}}\left(D_{\text{exp}}(t_i), D_{\text{sim}}(t_i)\right), \tag{S.14}$$

where

$$\hat{\mathcal{D}}_{\text{AD}}\left(D_{\text{exp}}(t_i), D_{\text{sim}}(t_i)\right) = -\tilde{N} - \sum_{j=1}^{\tilde{N}} \frac{2j-1}{\tilde{N}} \left[\ln\left(G_{\text{exp}}^{(i)}\left(d_{\text{sim}}^{(j)}(t_i)\right)\right) + \ln\left(1 - G_{\text{exp}}^{(i)}\left(d_{\text{sim}}^{(\tilde{N}+1-j)}(t_i)\right)\right)\right]. \tag{S.15}$$

In Eq. (S.15), $G_{\text{exp}}^{(i)}$ denotes the empirical cumulative distribution function for the experimental data at time $t_i$ and $d_{\text{sim}}^{(1)}(t_i), d_{\text{sim}}^{(2)}(t_i), \ldots, d_{\text{sim}}^{(\tilde{N})}(t_i)$ denote simulated values from the mathematical model in ascending order at time $t_i$, i.e. the elements of $D_{\text{sim}}(t_i)$. We follow [10] in Eq. (S.15) and avoid errors evaluating the natural logarithm when $P^{(j)}(t_i) = 0$ or $P^{(j)}(t_i) = 1$, corresponding to values outside of the range of data used to define $G_{\text{exp}}$, by computing

$$G_{\text{exp}}\left(d_{\text{sim}}^{(j)}(t_i)\right) \to \min\left(1 - 10^{-9}, \max\left(10^{-9}, G_{\text{exp}}\left(d_{\text{sim}}^{(j)}(t_i)\right)\right)\right). \tag{S.16}$$

In Supp. S2.1, we compare results obtained using this distance function based on the Anderson-Darling statistic to results obtained using distance functions based on the Cramer von Mises statistic and the Kolomogorov-Smirnov distance. For all of our choices of ABC distance functions a smaller distance corresponds to closer agreement between the experimentally measured data and the simulated data.

*ABC algorithm.* We use an established ABC Sequential Monte Carlo (SMC) algorithm, detailed in [10, 11], to estimate ABC posterior distributions for the statistical hyperparameters. The algorithm initially randomly generates 2000 parameter combinations sampled from uniform priors for $m_r$, $s_r$, $m_K$ and $s_K$. We refer to each of these 2000 parameter combinations as an ABC particle. Next, the algorithm computes the ABC distance for each of the ABC particles and orders the particles from the lowest ABC distance to the highest ABC distance. The 1500 (75%) ABC particles with the highest ABC distance, and so poorest agreement between the experimentally measured data and the simulated data, are then discarded. The first ABC error threshold, $\varepsilon$, is set equal to the highest remaining ABC distance. The 1500 replacement particles are then subsampled from the 500 remaining particles and perturbed using a multivariate Gaussian perturbation kernel. The covariance of this kernel is set equal to two times the empirical covariance of the retained ABC particles [11]. Following [10, 11], we allow the algorithm to proceed until a target acceptance probability is reached or until the maximum number of steps, set to 20, is reached. We also adapt the algorithm to terminate at a target ABC error threshold. We now explain when we use each terminating condition.

For each of the synthetic data studies with low, intermediate, and high $r$ (Fig. 6), we first use the ABC-SMC algorithm with a target acceptance probability set equal to 0.005. Next, we determine the ABC particle with the lowest ABC distance and recompute its ABC distance 1000 times. We perform this calculation 1000 times because the heterogeneous mathematical model is stochastic (due to sampling of noise from the cell-only experimental control data, particle-only control data, and sampling of the lognormal distributions). We then compute the median ABC distance, $\varepsilon_m$, from the 1000 distances for that particular ABC particle. We then report results in Fig. 6 using the ABC-SMC algorithm performed with a target ABC error threshold equal to the minimum $\varepsilon_m$ across the low, intermediate, and high $r$ data, which is 12.4 for the Anderson-Darling ABC distance function. We use this same ABC error threshold for the synthetic data study in Figs. 2-5. We use the same ABC error threshold across synthetic data studies in Fig. 2-6 to support a fair comparison of results across data sets.

For experimental data studies and for the comparison of results with different ABC distance functions, we first use the ABC-SMC algorithm with a target acceptance probability set equal to 0.005. We then determine the ABC particle with the lowest ABC distance, recompute its ABC distance 1000 times, and determine the median distance $\varepsilon_d$. Next, we use the ABC-SMC algorithm with a target ABC error threshold set equal to $\varepsilon_d$. This approach means that we may have a different ABC error threshold for each experimental data set and for each data set that we analyse with different ABC distance functions. We take this approach for computational efficiency so that the ABC-SMC algorithms terminate in a reasonable timeframe (24 hours).

*Prior distributions for statistical hyperparameters.* We assume uniform priors for all statistical hyperparameters throughout. In Table S3, we specify the uniform bounds that we use to generate results for each figure. Note that we use the same bounds for all experimental data, and we use narrower bounds to report the final results of the synthetic data studies to aid efficient inference.

For the analysis of optimal experimental designs, we choose the bounds for the uniform priors close to the known values used to generate the data to aid efficient inference (Fig. 9 in Table S3). In particular, we set the lower and upper bounds for $m_r$ and $s_r$ to be an order of magnitude below and above the pre-specified mean values, respectively. Furthermore, we set the uniform prior bounds to be $(5.0, 15.0)$ for $m_K$, and $(0.01, 4.0)$ for $s_K$, which are centred about and close to the respective pre-specified mean values of 10.0 and 2.0 .

| Fig. | Data | $m_r$ | $s_r$ | $m_K$ | $s_K$ |
|---|---|---|---|---|---|
| 7, 8(J-L), S5-S7 | Experimental | $(10^{-11}, 1)$ | $(10^{-11}, 1)$ | $(0.1, 100.0)$ | $(0.01, 200)$ |
| 2-5 | Synthetic | $(10^{-11}, 10^{-4})$ | $(10^{-11}, 10^{-4})$ | $(0.1, 100.0)$ | $(0.01, 100)$ |
| 6, 8(A-I), S4 | Synthetic | $(10^{-11}, 10^{-4})$ | $(10^{-11}, 10^{-4})$ | $(0.1, 40.0)$ | $(0.01, 20)$ |
| 9 | Synthetic | $(0.1m_r^*, 10m_r^*)$ | $(0.1s_r^*, 10s_r^*)$ | $(5.0, 15.0)$ | $(0.01, 4.0)$ |

**Table S3: Bounds for uniform priors.** Bounds reported for $m_r$ [m s$^{-1}$], $s_r$ [m s$^{-1}$] , $m_K$ [particles cell$^{-1}$], and $s_K$ [particles cell$^{-1}$]. To report bounds used in Fig. 9, we let $m_r^*$, and $s_r^*$ represent the pre-specified values of $m_r$ and $s_r$ for low, intermediate, and high $r$.

*Posterior distributions for statistical hyperparameters.* We compute the posterior distributions using all 2000 ABC particles from the final iteration of the ABC-SMC algorithm. We compute 95% highest posterior density intervals for each statistical hyperparameter using standard methods [13].

*Posterior predictions for flow cytometry experimental measurements, $D_{\text{exp}}$.* We first plot $D_{\text{exp}}(t_i)$ for $i = 1, 2, \ldots, 6$ as a histogram with a bin width equal to 20. We then obtain the 2000 ABC particles from the final iteration of the ABC-SMC algorithm. For each of these

ABC particles, we generate $D_{\text{sim}}$ using the heterogeneous mathematical model (Sec. 3). Note that $D_{\text{exp}} = \{D_{\text{exp}}(t_i) \mid i = 1, 2, \ldots, 6\}$ and $D_{\text{sim}} = \{D_{\text{sim}}(t_i) \mid i = 1, 2, \ldots, 6\}$ are directly comparable by definition. We then generate 95% pointwise prediction intervals for each $t_i$ by estimating the 2.5% and 97.5% quantiles of the simulated data, $D_{\text{sim}}(t_i)$, at each midpoint of the histogram bins. The solid line displayed in the prediction interval corresponds to the median of the simulated data.

*Inferred distributions for $r$ and $K$.* To generate an inferred distribution for $r$, we first obtain 2000 samples of $m_r$ and $s_r$ from the 2000 ABC particles from the final iteration of the ABC-SMC algorithm. For each of these samples we simulate the lognormal distribution for $r$. We then generate 95% pointwise prediction intervals by estimating the 2.5% and 97.5% quantiles of the 2000 simulated lognormal distributions at different values of $r$. The solid line displayed in the prediction interval corresponds to the median of the simulated data. We generate an inferred distribution for $K$ similarly.

*Predictions for particles per cell, $P(t)$.* For 2000 samples of $m_r$ and $s_r$ from the 2000 ABC particles from the final iteration of the ABC-SMC algorithm, we simulate $P^{(j)}(t)$ for $0 \leq t \leq 24$ [hr] and $j = 1, 2, \ldots, \tilde{N}$. per Eq. (5). We then estimate 95% (50%) pointwise prediction intervals by estimating the 2.5% (25%) and 97.5% (75%) quantiles of the simulated data in steps of 0.5 [hr]. The solid line displayed in the prediction interval corresponds to the median of the simulated data.

*Predictions for percentage of cells close to $K$ at time $t$.* To explain only this prediction we vary notation. We let $P_i^{(j)}\left(t; r_i^{(j)}, K_i^{(j)}\right)$ represent the number of particles in simulated cell $j$ for ABC particle $i$ and let $r_i^{(j)}$ and $K_i^{(j)}$ represent the sampled values of $r$ and $K$ that characterise simulated cell $j$ for ABC particle $i$. We follow the method immediately above to generate these predictions for all 2000 ABC particles from the final iteration of the ABC-SMC algorithm. We then define the percentage of cells *close* to $K$ at time $t$ to be the percentage of simulated cells that have more associated particles than the $\beta$ fraction of their carrying capacity,

$$100 \times \frac{1}{2000} \frac{1}{\tilde{N}} \sum_{i=1}^{2000} \sum_{j=1}^{\tilde{N}} \mathbb{1}\left(P_i^{(j)}(t; r_i^{(j)}, K_i^{(j)}) - \beta K_i^{(j)}\right), \tag{S.17}$$

where $0 \leq \beta \leq 1$ and $\mathbb{1}(x)$ is an indicator function defined to be equal to 1 when $x > 0$ and equal to 0 when $x \leq 0$. The summation over $j$ indicates that we analyse all $\tilde{N}$ simulated cells for each ABC particle $i$, and the summation over $i$ indicates that we analyse all ABC particles in the ABC posterior. The term $1/(2000\tilde{N})$ is a normalising constant for the summations, and we multiply by 100 to convert the fraction to a percentage. In Fig. 5(B) we present results for $\beta = 0.50, 0.95, 0.99$.

#### S1.5 Parameter estimation for the homogeneous mathematical model

In Sec. 4.5, we use the method of least squares to connect the homogeneous mathematical model (Sec. 3.1.1) to experimental estimates of the number of particles per cell at each time point, $P_o(t_i)$ [particles cell$^{-1}$] for $t_i = 1, 2, 4, 8, 16, 24$ [hr] (defined from flow cytometry data per Eq. (10)) with initial condition $P_o(0) = 0$ [particles cell$^{-1}$]. Here, we present further details for this method of least squares approach.

Recalling that the homogeneous mathematical model is characterised by $r$ and $K$, we let $P(t_i; r, K)$ for $t_i = 1, 2, 4, 8, 16, 24$ [hr] denote the solution of the homogeneous mathematical model at time $t_i$ evaluated with known values of $r$ and $K$. We then formulate a least squares estimation problem,

$$E(r, K) = \sum_{i=1}^{I} (P_o(t_i) - P(t_i; r, K))^2, \tag{S.18}$$

Minimising Eq. (S.18), which corresponds to minimising the sum of squared residuals, determines the best-fit point-estimates $\hat{r}$ and $\hat{K}$. We perform this minimisation numerically using the Nelder-Mead local optimisation routine, with default stopping criteria, within the NLopt.jl Julia optimisation package [12]. The *best-fit* point-estimate prediction for the number of particles per cell then corresponds to the solution of the homogeneous mathematical model evaluated at the best-fit point-estimates $\hat{r}$ and $\hat{K}$, namely $P(t; \hat{r}, \hat{K})$.

### S2 Results

#### S2.1 Different ABC distance functions and nano-engineered particles

In the main manuscript, we present results using an ABC distance function based on the Anderson-Darling statistic. Here, we compare these results to those obtained using ABC distance functions based on the Cramer von Mises statistic and the Kolmogorov-Smirnov distance. We present these results for the synthetic data discussed in Fig. 6, noting that results for the other synthetic data studies are very similar. We also present results for the experimental data discussed in Fig. 7, namely the 150 nm PMA-core-shell particles, and for two additional experimental data sets that are not discussed in the main manuscript, namely the 214 nm PMA-capsules and 633 nm PMA core-shell particles.

We first define the three ABC distance functions that we consider:

- *Anderson-Darling.* We define the ABC distance function based on the Anderson-Darling statistic in Eqs. (S.14) and (S.15).

- *Cramer von Mises.* We define the ABC distance function based on the Cramer von Mises statistic to be,

$$\mathcal{D}_{\text{CVM}}(D_{\text{exp}}, D_{\text{sim}}) = \sum_{i=1}^{6} \hat{\mathcal{D}}_{\text{CVM}}(D_{\text{exp}}(t_i), D_{\text{sim}}(t_i)), \tag{S.19}$$

  where

$$\hat{\mathcal{D}}_{\text{CVM}}(D_{\text{exp}}(t_i), D_{\text{sim}}(t_i)) = \frac{1}{12\tilde{N}} \sum_{j=1}^{\tilde{N}} \left[ \frac{2j-1}{2\tilde{N}} - G_{\text{exp}}^{(i)}\left(d_{\text{sim}}^{(j)}(t_i)\right) \right]^2. \tag{S.20}$$

  In Eq. (S.20), $G_{\text{exp}}^{(i)}$ denotes the empirical cumulative distribution function for the experimental data at time $t_i$ and $d_{\text{sim}}^{(1)}(t_i), d_{\text{sim}}^{(2)}(t_i), \ldots, d_{\text{sim}}^{(\tilde{N})}(t_i)$ denote simulated values from the mathematical model in ascending order at time $t_i$, i.e. the elements of $D_{\text{sim}}(t_i)$.

- *Kolmogorov Smirnov.* We define the ABC distance function based on the Kolmogorov-Smirnov distance to be,

$$\mathcal{D}_{\text{KS}}(D_{\text{exp}}, D_{\text{sim}}) = \sum_{i=1}^{6} \hat{\mathcal{D}}_{\text{KS}}(D_{\text{exp}}(t_i), D_{\text{sim}}(t_i)), \tag{S.21}$$

  where

$$\hat{\mathcal{D}}_{\text{KS}}(D_{\text{exp}}(t_i), D_{\text{sim}}(t_i)) = \max_{j=1,2,\ldots,2\tilde{N}} \left| G_{\text{sim}}^{(i)}(x_j) - G_{\text{exp}}^{(i)}(x_j) \right|. \tag{S.22}$$

  In Eq. (S.22), $G_{\text{exp}}^{(i)}$ and $G_{\text{sim}}^{(i)}$ denote the empirical cumulative distribution functions for the experimental and simulated data at time $t_i$, respectively. Furthermore, the $x_j$ for $j = 1, 2, \ldots, 2\tilde{N}$ represent the $\tilde{N}$ elements of $D_{\text{exp}}(t_i)$ and the $\tilde{N}$ elements $D_{\text{sim}}(t_i)$.

We now compare results obtained using the three ABC distance functions for the synthetic data with intermediate $r$ first discussed in Fig. 6. Posterior distributions for all statistical hyperparameters demonstrate excellent agreement across all three ABC distance functions and capture the known parameters used to generate the data (Fig. S4(A-D)). Consequently, all posterior predictions across the three ABC distance functions also demonstrate excellent agreement. This includes predictions for fluorescence data (Fig. S4(E-J)), the median of the inferred distributions for $r$ and $K$ (Fig. S4(K,L)), and predictions for $P(t)$ (Fig. S4(M)).

We next compute the Anderson-Darling ABC distance for the ABC samples generated using the Anderson-Darling, Cramer von Mises, and Kolmogorov Smirnov ABC distance functions (Fig. S4(N)). These distances are similar suggesting that our choice of ABC algorithm and thresholds allow for a fair comparison of the ABC distance functions. We observe similar results for the Cramer von Mises and Kolmogorov Smirnov ABC distances (Fig. S4(O,P)).

These synthetic data results in Fig. (S4) suggest that, in the absence of model misspecification, estimates of the statistical hyperparameters and subsequent predictions are largely independent of whether the Anderson-Darling, Cramer von Mises, or Kolmogorov Smirnov ABC distance function is used. Results for the other synthetic data studies that we consider in Figs. (2-6) are similar. Other ABC distance functions can also be explored using our methods.

We now consider the 150 nm experimental data first discussed in Fig. 7 with the three ABC distance functions. These posterior distributions demonstrate good agreement across the ABC distance functions (Fig. S5(A-D)). In particular, the posterior distributions for each statistical hyperparameter across the ABC distance functions overlap and are of the same order of magnitude. These differences in the posterior distributions result in small differences in predictions for fluorescence data (Fig. S5(E-J)). Furthermore, the median of the inferred distributions for $r$ for each ABC distance function demonstrates excellent agreement in general with small differences close to zero (Fig. S5(K)). The median of the inferred distributions for $r$ for each of the ABC distance functions also demonstrates good agreement(Fig. S5(L)). Predictions for $P(t)$ from each of the ABC distance functions demonstrate excellent agreement at the median and 50% intervals. However, there is some variability in the upper tails of the predictions, notably at the upper boundary of the 95% prediction interval. As we demonstrate that our ABC methods accurately recover known parameters for the synthetic data in the absence of model misspecification, we attribute these differences for the experimental data to model misspecification. This model misspecification also results in additional variability in the ABC distances for the ABC samples generated using the three different ABC distance functions in comparison to the synthetic data studies.

Repeating this analysis for the 214 nm experimental data (Fig. S6) and 633 nm experimental data (Fig. S7), we obtain similar results regarding the differences that may arise across the three ABC distance functions. We do not make direct comparisons between results obtained for the different experimental data sets as we use different ABC error thresholds to generate these results.

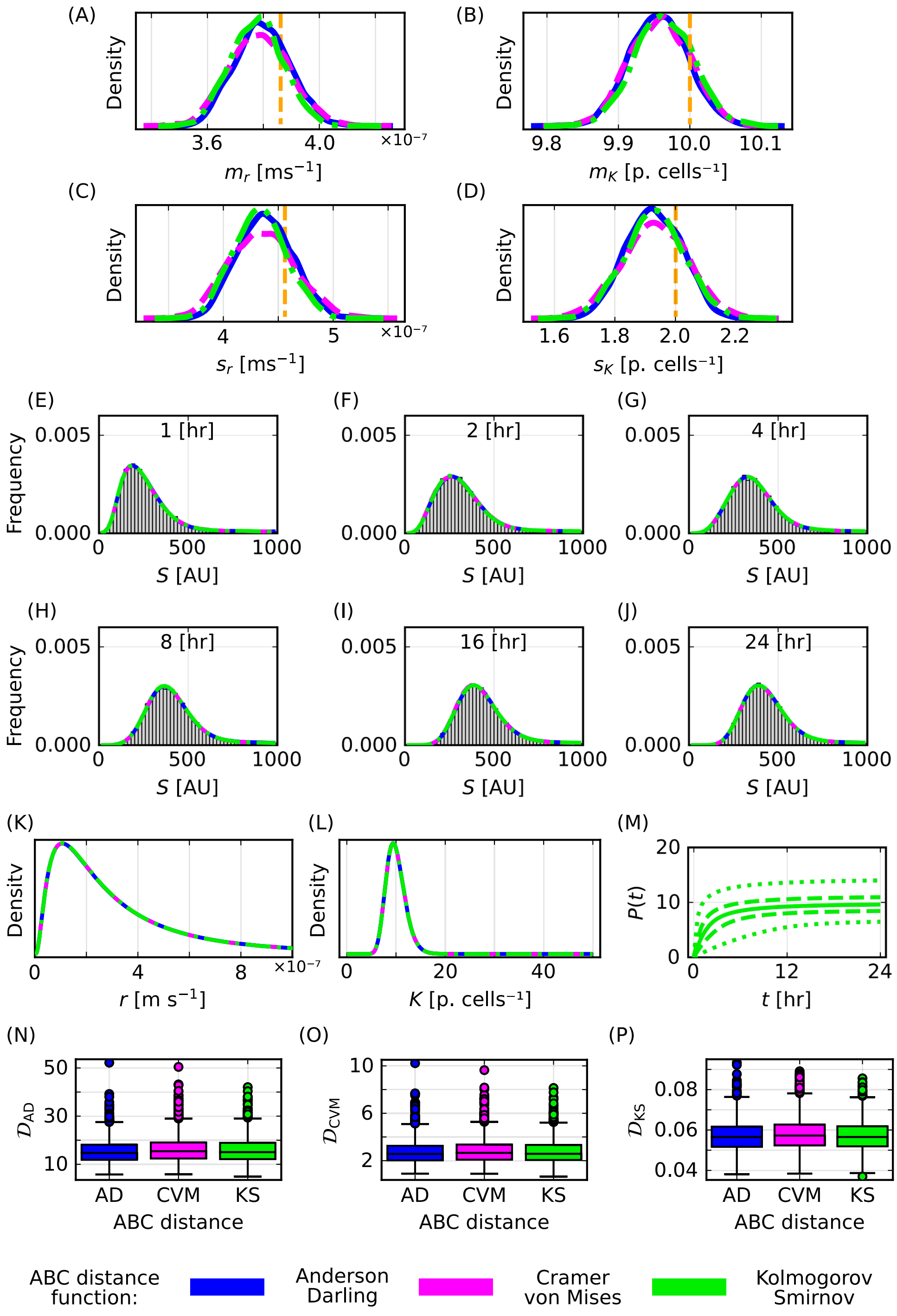

**Figure S4: Results for synthetic data using different ABC distance functions.** (A-D) Posterior distributions for the statistical hyperparameters (A) $m_r$ [m s$^{-1}$], (B) $m_K$ [particles cell$^{-1}$], (C) $s_r$ [m s$^{-1}$], and (D) $s_K$ [particles cell$^{-1}$] with known parameters (vertical orange). (E-J) Histograms of synthetic fluorescence data (grey) and median predictions for the heights of histogram bars from the mathematical model for $t = 1, 2, 4, 8, 16, 24$ [hours]. (K)-(L) Median inferred distribution for (K) $r$ [m s$^{-1}$] and (L) $K$ [particles cell$^{-1}$]. (M) Predictions of $P(t)$ [particles cell$^{-1}$] with median (dotted), 50% prediction interval (dash-dotted), and 95% prediction interval (solid). (O) Anderson-Darling distance of ABC samples obtained using the ABC-SMC algorithm with the Anderson-Darling, Cramer von Mises, and Kolmogorov Smirnov distance functions. (N) Cramer von Mises distance and (P) Kolmogorov Smirnov distance of ABC samples described in (O). Colours represent the Anderson-Darling (blue), Cramer von Mises (magenta), and Kolomogorov-Smirnov (green) ABC distance functions.

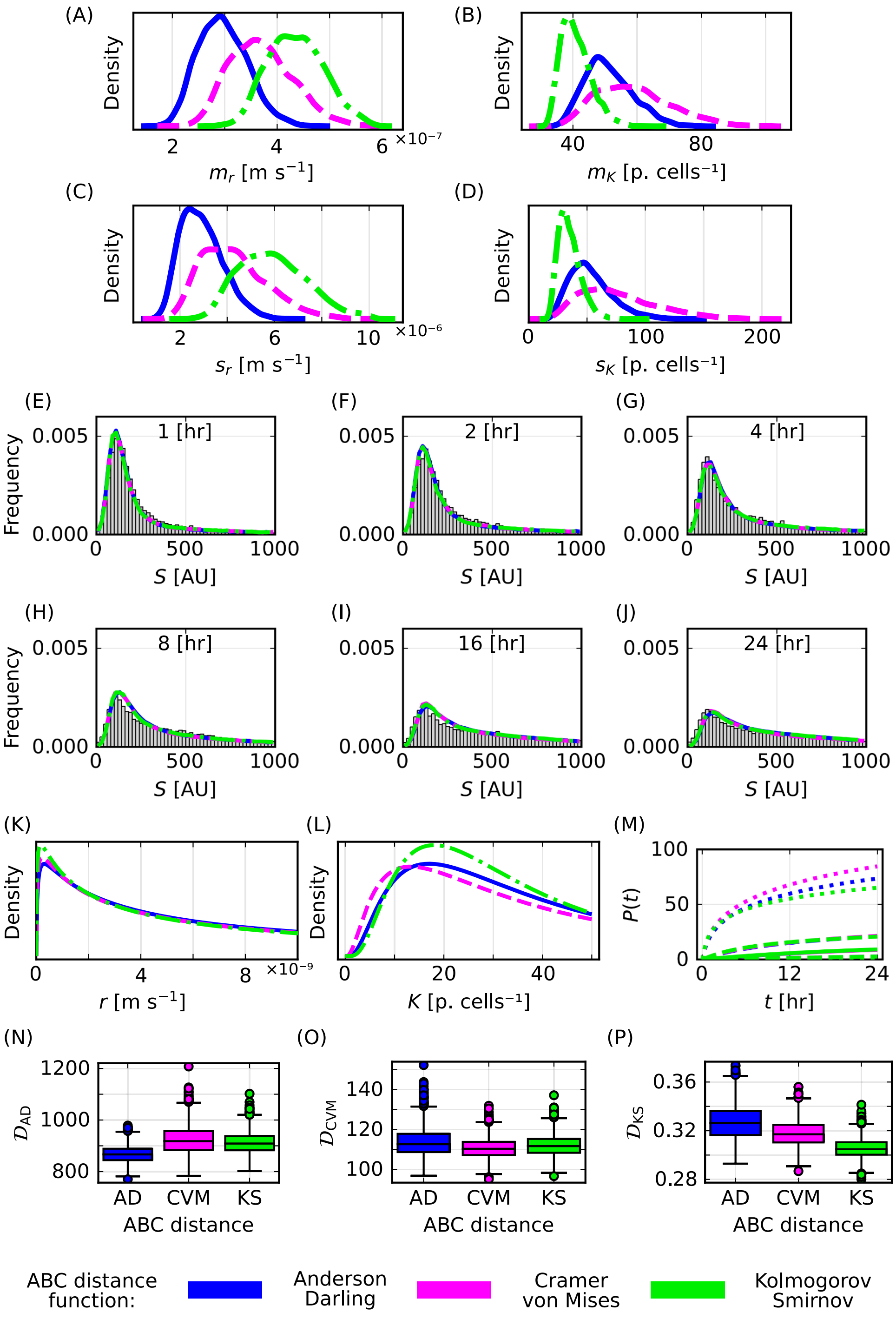

**Figure S5: Results for 150 nm PMA-core-shell particles experimental data using different ABC distance functions.** (A-D) Posterior distributions for the statistical hyper-parameters (A) $m_r$ [m s$^{-1}$], (B) $m_K$ [particles cell$^{-1}$], (C) $s_r$ [m s$^{-1}$], and (D) $s_K$ [particles cell$^{-1}$]. (E-J) Histograms of experimental fluorescence data (grey) and median predictions for the heights of histogram bars from the mathematical model for $t = 1, 2, 4, 8, 16, 24$ [hours]. (K)-(L) Median inferred distribution for (K) $r$ [m s$^{-1}$] and (L) $K$ [particles cell$^{-1}$]. (M) Predictions of $P(t)$ [particles cell$^{-1}$] with median (dotted), 50% prediction interval (dash-dotted), and 95% prediction interval (solid). (O) Anderson-Darling distance of ABC samples obtained using the Anderson-Darling, Cramer von Mises, and Kolmogorov Smirnov distance functions. (N) Cramer von Mises distance and (P) Kolmogorov Smirnov distance of ABC samples described in (O). Colours represent the Anderson-Darling (blue), Cramer von Mises (magenta), and Kolomogorov-Smirnov (green) ABC distance functions.

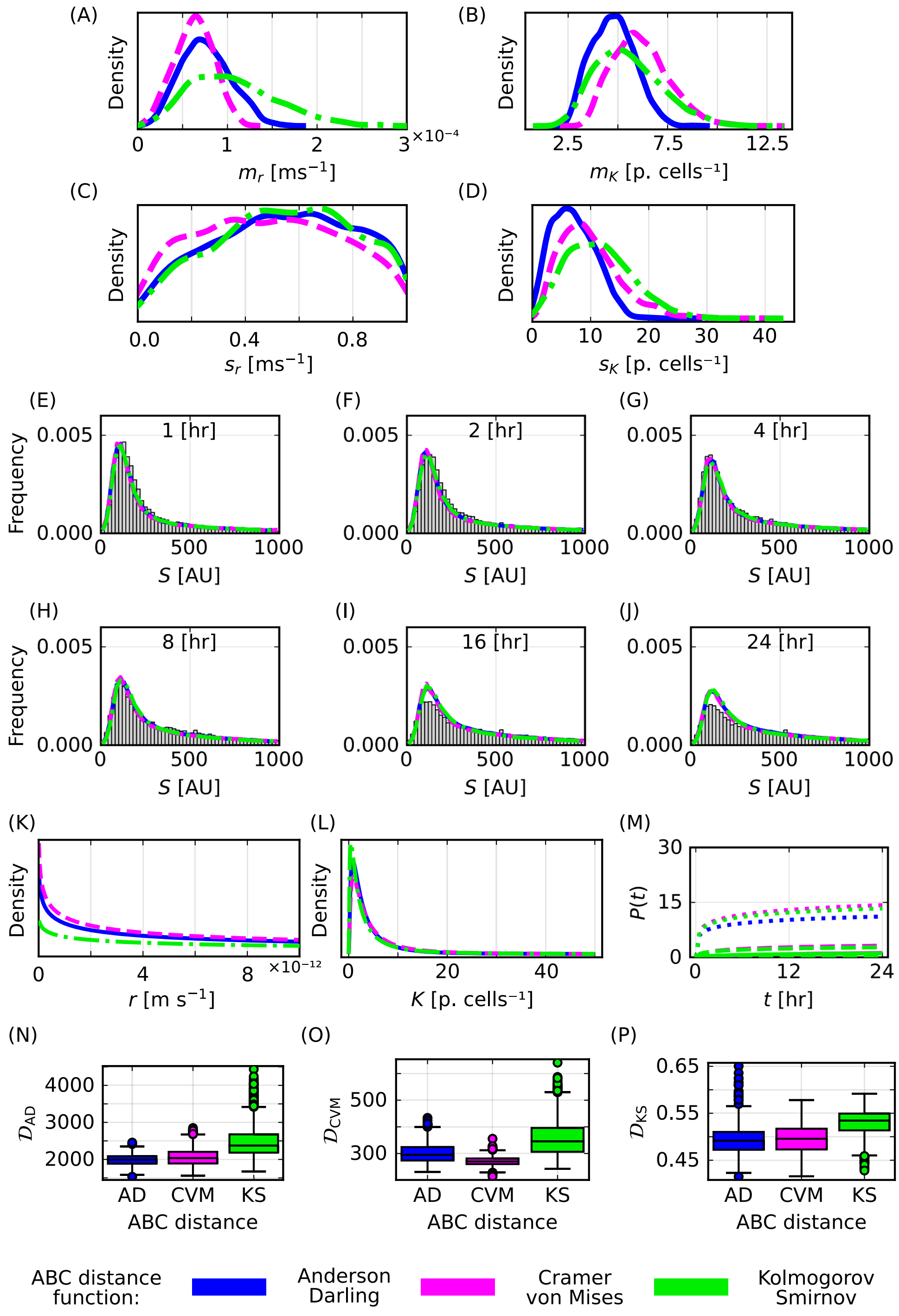

**Figure S6: Results for 214 nm PMA-capsules experimental data using different ABC distance functions.** (A-D) Posterior distributions for the statistical hyperparameters (A) $m_r$ [m s$^{-1}$], (B) $m_K$ [particles cell$^{-1}$], (C) $s_r$ [m s$^{-1}$], and (D) $s_K$ [particles cell$^{-1}$]. (E-J) Histograms of experimental fluorescence data (grey) and median predictions for the heights of histogram bars from the mathematical model for $t$ = 1, 2, 4, 8, 16, 24 [hours]. (K)-(L) Median inferred distribution for (K) $r$ [m s$^{-1}$] and (L) $K$ [particles cell$^{-1}$]. (M) Predictions of $P(t)$ [particles cell$^{-1}$] with median (dotted), 50% prediction interval (dash-dotted), and 95% prediction interval (solid). (O) Anderson-Darling distance of ABC samples obtained using the Anderson-Darling, Cramer von Mises, and Kolmogorov Smirnov distance functions. (N) Cramer von Mises distance and (P) Kolmogorov Smirnov distance of ABC samples described in (O). Colours represent the Anderson-Darling (blue), Cramer von Mises (magenta), and Kolomogorov-Smirnov (green) ABC distance functions.

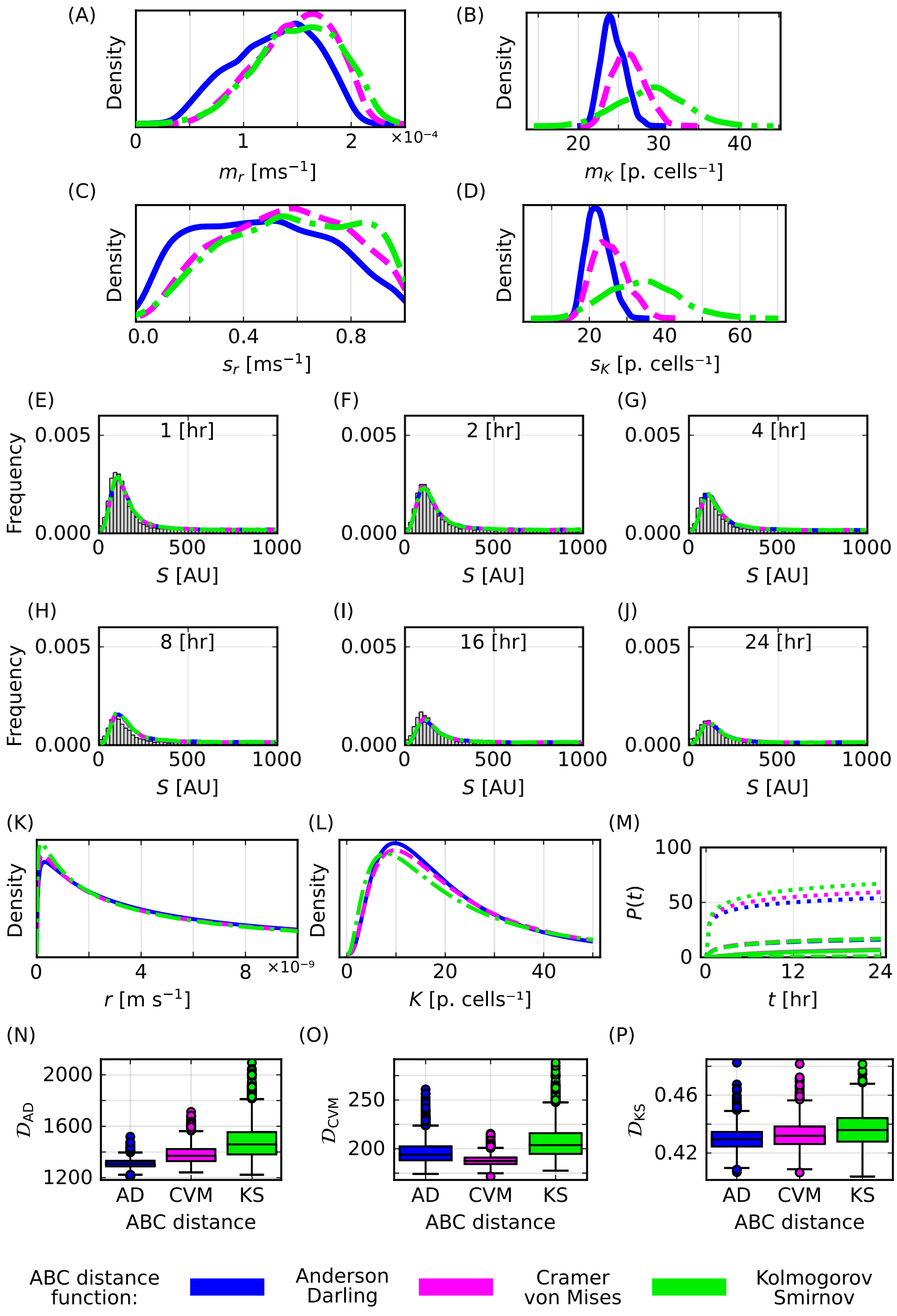

**Figure S7: Results for 633 nm PMA core-shell particles experimental data using different ABC distance functions.** (A-D) Posterior distributions for the statistical hyperparameters (A) $m_r$ [m s$^{-1}$], (B) $m_K$ [particles cell$^{-1}$], (C) $s_r$ [m s$^{-1}$], and (D) $s_K$ [particles cell$^{-1}$]. (E-J) Histograms of experimental fluorescence data (grey) and median predictions for the heights of histogram bars from the mathematical model for $t$ = 1, 2, 4, 8, 16, 24 [hours]. (K)-(L) Median inferred distribution for (K) $r$ [m s$^{-1}$] and (L) $K$ [particles cell$^{-1}$]. (M) Predictions of $P(t)$ [particles cell$^{-1}$] with median (dotted), 50% prediction interval (dash-dotted), and 95% prediction interval (solid). (O) Anderson-Darling distance of ABC samples obtained using the Anderson-Darling, Cramer von Mises, and Kolmogorov Smirnov distance functions. (N) Cramer von Mises distance and (P) Kolmogorov Smirnov distance of ABC samples described in (O). Colours represent the Anderson-Darling (blue), Cramer von Mises (magenta), and Kolomogorov-Smirnov (green) ABC distance functions.

#### S2.2 Regimes with parameter identifiability challenges

In Sec. 4.3 we explore parameter identifiability challenges for low, intermediate, and high $r$. In Fig. S8 we present posterior predictions for this synthetic flow cytometry data. All predictions demonstrate excellent agreement with the data. Note that the classifications of low, intermediate, and high $r$ are based on predictions of $P(t)$ from mathematical modelling and are relative to $K$.

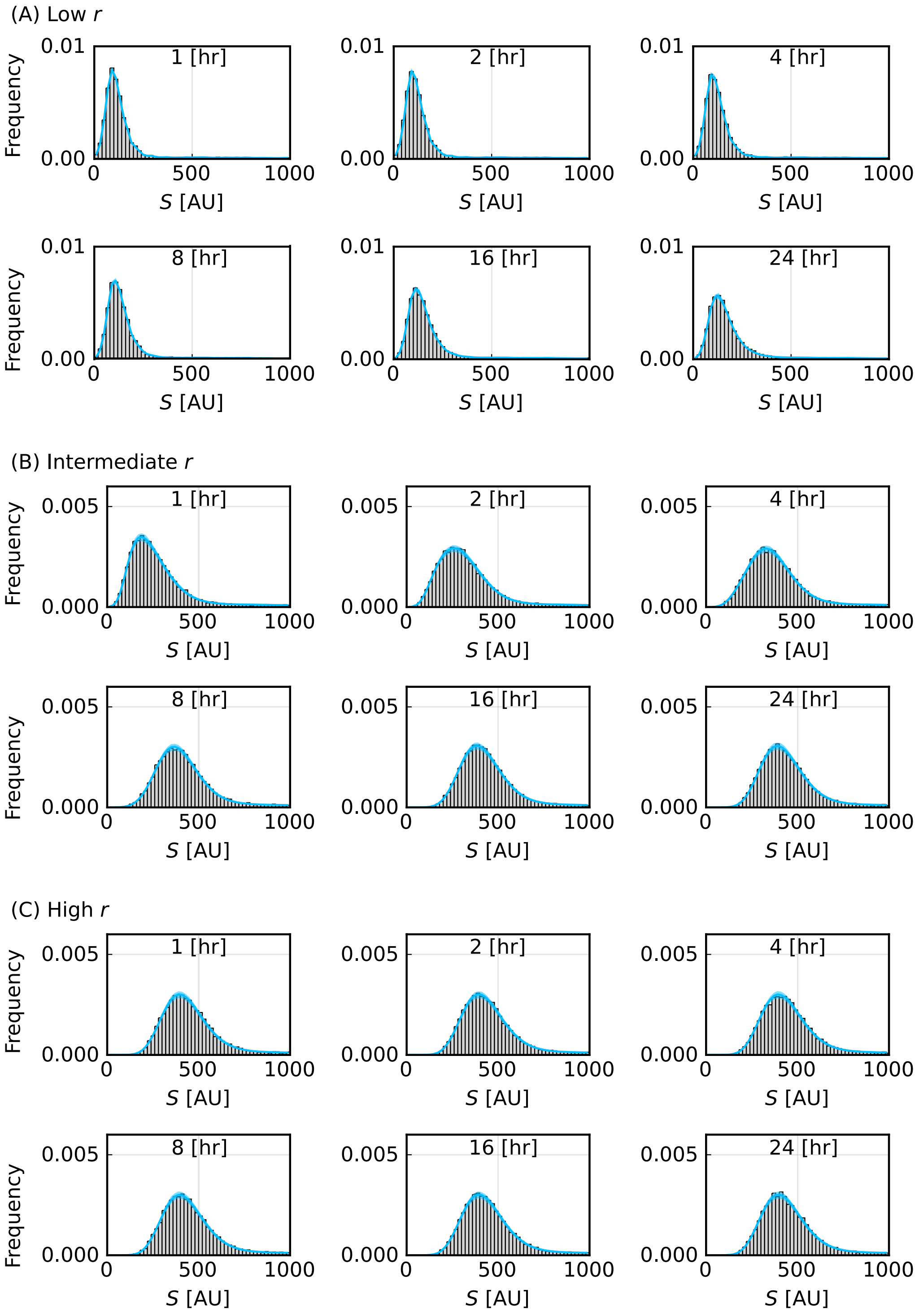

**Figure S8: Posterior predictions for synthetic flow cytometry data generated with low, intermediate, and high $r$.** Results for (A) low $r$, (B) intermediate $r$, and (C) high $r$.

#### S2.3 Comparison to previous methods: Role of control data

In Sec. 4.5, we compare our new method for predicting the number of particles per cell, $P(t)$, to a previous method that assumes a homogeneous cell population and obtains point estimates using the method of least squares. In Fig. 8(A-I), we show that for synthetic data generated using the heterogeneous mathematical model the least squares approach systematically overestimates the median of known values of $P(t)$. Here, we show that the least squares approach also systematically overestimates the median of known values of $P(t)$ for synthetic data generated using the homogeneous model (Fig. S9A-C). We then show that if the cell-only and particle-only control data are normally distributed instead of right-skewed, then the least squares approach accurately estimates the median of known values of $P(t)$ for both the homogeneous and heterogeneous mathematical models (Figs. S9D-F, S10). These results suggest that the systematic overestimation of $P(t)$ from the least squares approach is a consequence of the cell-only and particle-only control experimental data being right-skewed.

To generate the synthetic data in Fig. S9, we assume that the homogeneous mathematical model is valid and that there is no variability in $r$ and $K$. We then simulate the homogeneous mathematical model with known values of $r$ and $K$ (set equal to same known values of $m_r$ and $m_K$ as in Fig. 8) and determine the known values of $P(t)$. We convert these known values of $P(t)$ to synthetic flow cytometry data following Sec. 3.2. This means that we sample from the right-skewed experimental control data, $D_{\text{cells}}$ and $D_{\text{particles}}$. Following Sec. 4.5, we estimate $P_{\text{o}}(t_i)$ using the data transformation in Eq. (10) and fit to these $P_{\text{o}}(t_i)$ using the method of least squares. In Fig. S9(A-C), we observe that the least squares approach systematically overestimates the median of known values of $P(t)$ from synthetic data.

In Fig. S9(D-F), we re-perform this analysis with new synthetic control data, $D^{\text{syn}}_{\text{cells}}$ and $D^{\text{syn}}_{\text{particles}}$, replacing the experimentally measured control data $D_{\text{cells}}$ and $D_{\text{particles}}$. To facilitate a fair comparison, we base our new synthetic control data on the experimentally measured control data, $D_{\text{cells}}$ and $D_{\text{particles}}$, but importantly now assume that the synthetic control data are normally distributed with zero skew. In particular, we generate these data by sampling from normal distributions,

$$
\begin{aligned}
D^{\text{syn}}_{\text{cells}} &= \{d^{(j)}_{\text{cells}} \mid d^{(j)}_{\text{cells}} \sim N\left(m_{\text{cells}}, s^2_{\text{cells}}\right) \text{ for } j = 1, 2, \ldots, |D_{\text{cells}}|\}, \\
D^{\text{syn}}_{\text{particles}} &= \{d^{(j)}_{\text{particles}} \mid d^{(j)}_{\text{particles}} \sim N\left(m_{\text{particles}}, s^2_{\text{particles}}\right) \text{ for } j = 1, 2, \ldots, |D_{\text{particles}}|\},
\end{aligned}
\tag{S.23}
$$

where $N(m, s^2)$ represents a normal distribution with mean $m$ and standard deviation $s$, $m_{\text{cells}}$ and $s_{\text{cells}}$ represent the mean and standard deviation of $D_{\text{cells}}$, $m_{\text{particles}}$ and $s_{\text{particles}}$ represent the mean and standard deviation of $D_{\text{particles}}$, and we let $|D_{\text{cells}}|$ and $|D_{\text{particles}}|$ denote the number of elements of $D_{\text{cells}}$ and $D_{\text{cells}}$, respectively. In Fig. S9(D-F), we observe that the least squares approach now generates estimates of $P(t)$ that agree with the median of known values of $P(t)$ from synthetic data.

For the heterogeneous model, we re-perform the analysis in Fig. 8(A-I) with the new synthetic control data $D^{\text{syn}}_{\text{cells}}$ and $D^{\text{syn}}_{\text{particles}}$ (Eq. (S.23)) replacing the experimentally measured control data $D_{\text{cells}}$ and $D_{\text{particles}}$. We observe that the estimates of $P(t)$ from the least squares approach now agree with the median of known values of $P(t)$ from synthetic data (Fig. S10).

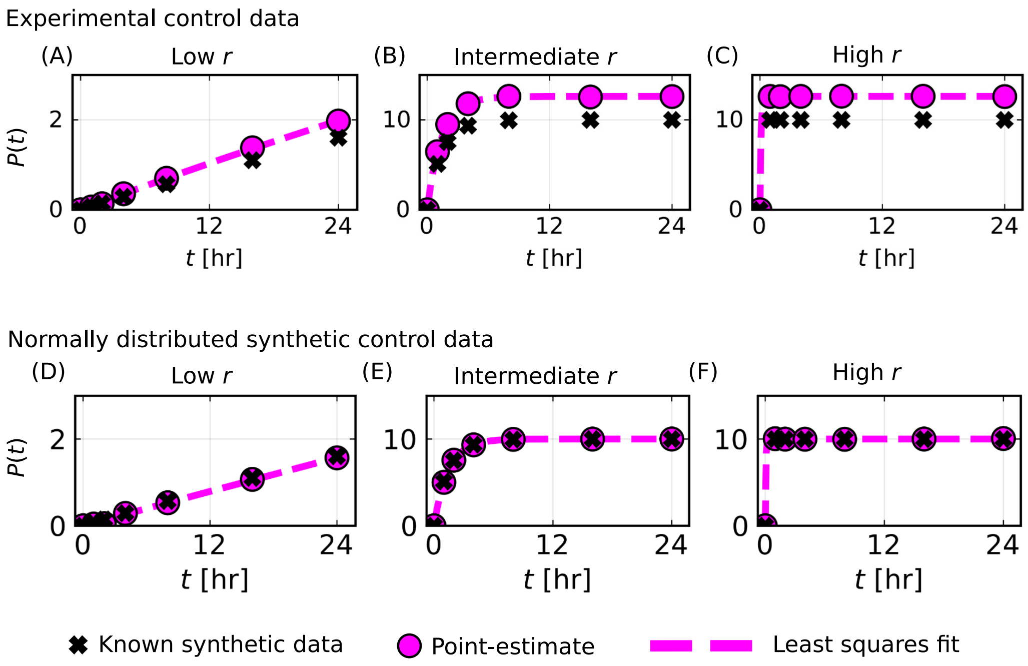

**Figure S9: Comparison of results for the point-estimate method applied to synthetic data generated using the homogeneous model and two types of control data.** Results in (A-C) are for experimentally measured control data. Results in (D-F) are for synthetic normally distributed control data. (A-F) $P_o(t_i)$ [particles cell$^{-1}$] from Eq. (10) (magenta circles) and solution of the homogeneous cell population mathematical model evaluated at the best-fit parameter values for $r$ and $K$ from the method of least squares (magenta dashed) compared to known synthetic data for $P(t)$ [particles cell$^{-1}$] (black crosses).

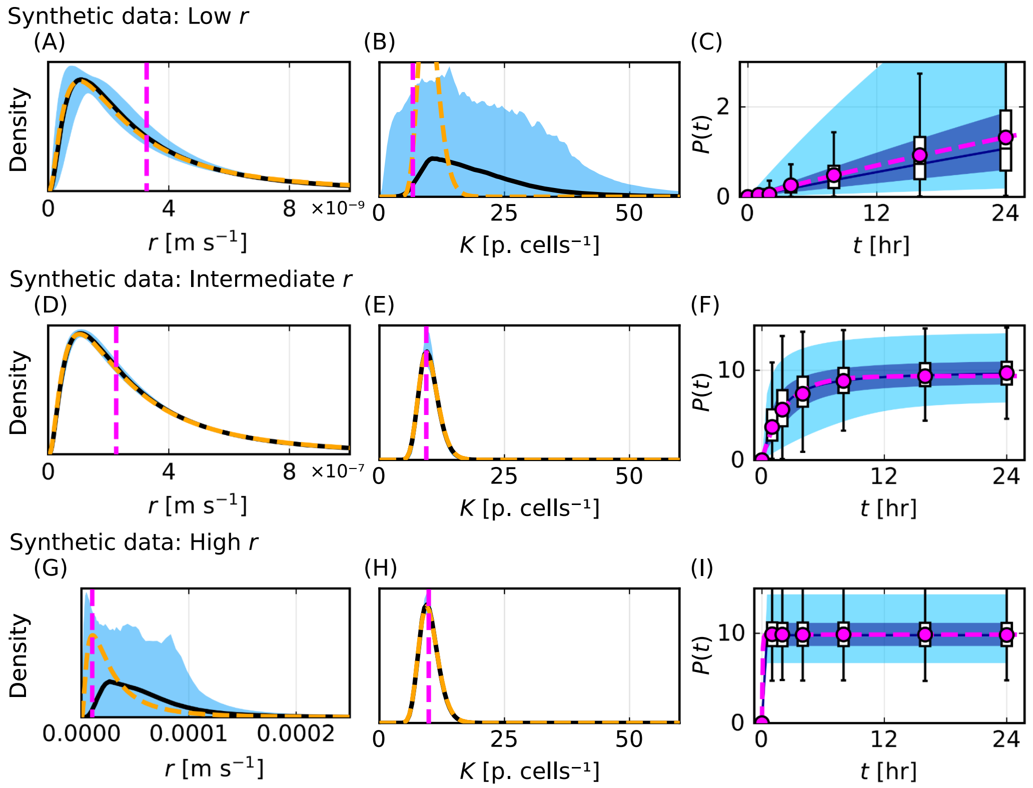

**Figure S10: Comparison of point estimate method with inferred distributions and predictions obtained using our new method for synthetic and normally distributed $D_{\text{cells}}$ and $D_{\text{particles}}$.** (A,D,G) Best-fit parameter values for $r$ from the homogeneous model with the method of least squares (magenta vertical dashed) compared to inferred distribution for $r$ from our new method. (B,E,H) Best-fit parameter values for $K$ from the homogeneous model with the method of least squares (magenta vertical dashed) compared to inferred distribution for $r$ from our new method. (C,F,I) $P_{\text{o}}(t_i)$ [particles cell$^{-1}$] from Eq. (10) (magenta circles) and solution of the homogeneous cell population mathematical model evaluated at the best-fit parameter values for $r$ and $K$ from the method of least squares (magenta dashed) compared to prediction of $P(t)$ [particles cell$^{-1}$] from our new method.

#### S2.4 Optimal experimental time points

In Table. S4 we present the time points that define the nine selected designs that we present in Fig. 9(A), their normalised utility for each particle-cell scenario, their rankings for each particle-cell scenario, their average ranking across the three particle-cell scenarios, and an overall ranking that is based on the average ranking across the three particle-cell scenarios.

| Design | Time point | | | | | | Normalised mean utility | | | Ranking | | | | Average ranking all data |
|---|---|---|---|---|---|---|---|---|---|---|---|---|---|---|
| | $t_1$ | $t_2$ | $t_3$ | $t_4$ | $t_5$ | $t_6$ | Low $r$ | Int $r$ | High $r$ | Low $r$ | Int $r$ | High $r$ | Overall | |
| Early | 0.5 | 1 | 2 | 4 | 6 | 8 | 0.014 | 0.210 | 0.922 | 3003 | 2608 | 1251 | **2866** | 2287 |
| Middle | 6 | 8 | 10 | 12 | 14 | 16 | 0.239 | 0.032 | 0.558 | 2333 | 2928 | 2943 | **3002** | 2735 |
| Late | 14 | 16 | 18 | 20 | 22 | 24 | 0.660 | 0.003 | 0.492 | 244 | 3003 | 3003 | **2685** | 2083 |
| Uniform | 4 | 8 | 12 | 16 | 20 | 24 | 0.743 | 0.197 | 0.638 | 135 | 2640 | 2594 | **2119** | 1790 |
| Experimental data | 1 | 2 | 4 | 8 | 16 | 24 | 0.320 | 0.649 | 0.781 | 1808 | 565 | 1942 | **1430** | 1438 |
| Optimal low $r$ | 8 | 10 | 14 | 20 | 22 | 24 | 1.000 | 0.039 | 0.573 | 1 | 2922 | 2914 | **2430** | 1946 |
| Optimal int $r$ | 0.5 | 1 | 2 | 20 | 22 | 24 | 0.204 | 1.000 | 0.937 | 2516 | 1 | 1064 | **855** | 1194 |
| Optimal high $r$ | 0.5 | 4 | 6 | 12 | 16 | 20 | 0.271 | 0.449 | 1.000 | 2119 | 1652 | 1 | **1000** | 1257 |
| Optimal unknown $r$ | 0.5 | 4 | 12 | 14 | 22 | 24 | 0.633 | 0.568 | 0.994 | 298 | 939 | 16 | **1** | 418 |

**Table S4: Time points, normalised mean utility, and rankings for nine select designs.**
